# The U1 snRNP component RBP45d regulates temperature-responsive flowering in *Arabidopsis thaliana*

**DOI:** 10.1101/2021.06.14.448173

**Authors:** Ping Chang, Hsin-Yu Hsieh, Shih-Long Tu

## Abstract

Pre-mRNA splicing is a crucial step of gene expression whereby the spliceosome produces constitutively and alternatively spliced transcripts that not only diversify the transcriptome but also play essential functions during plant development and responses to environmental changes. Numerous evidences indicate that regulation at the pre-mRNA splicing step is important for flowering time control, however the components and detailed mechanism underlying this process remain largely unknown. Here, we identified a previously unknown splicing factor in *Arabidopsis thaliana*, RNA BINDING PROTEIN 45d (RBP45d), a member of the RBP45/47 family. Using sequence comparison and biochemical analysis, we determined that RBP45d is a component of the U1 small nuclear ribonucleoprotein (U1 snRNP) with functions distinct from other family members. RBP45d associates with the U1 snRNP by interacting with pre-mRNA-processing factor 39a (PRP39a) and directly regulates alternative splicing (AS) for a specific set of genes. Plants with loss of RBP45d function exhibit defects in temperature-induced flowering potentially due to the mis-regulation of temperature-sensitive AS by RBP45d and PRP39a of the key flowering gene *FLOWERING LOCUS M*. Taken together, we report that RBP45d is a novel U1 snRNP component in plants that functions together with PRP39a in temperature-mediated flowering.

## INTRODUCTION

Eukaryotic genes are interrupted by introns, the intervening sequences between exons that need to be removed after being transcribed into precursor messenger RNAs (pre-mRNAs). The removal of intronic regions from pre-mRNAs is carried out by a dynamic macromolecular protein–RNA complex, the spliceosome, which is composed of U1, U2, U4/U6 and U5 small nuclear ribonucleoproteins (snRNPs) (Will and Lührmann, 2011). Pre-mRNA splicing starts with the recognition of the 5’- and 3’-splice sites (SS), followed by two transesterification steps: branching and 3’-SS ligation. The 5’-SS is recognized through base pairing with U1 snRNA, with the help of U1 snRNP (Lerner et al., 1980; Zhuang and Weiner, 1986; Verhage et al., 2017; Plaschka et al., 2018), while the 3’-SS is recognized together with the branching point by SPLICING FACTOR 1 (SF1) and U2 snRNP auxiliary factors (U2AF35 and U2AF65) (Abovich and Rosbash, 1997). However, sometimes splice site selection is altered and results in the generation of two or more mRNA isoforms from the same pre-mRNA, a phenomenon that is termed alternative splicing (AS). AS is a widespread mechanism that greatly increases the complexity of the transcriptome and proteome, with ∼60% of genes in Arabidopsis (*Arabidopsis thaliana*) and ∼95% genes in human undergoing AS (Pan et al., 2008; Marquez et al., 2012). In addition to creating a more complex proteome, AS often generates mRNA isoforms possessing pre-mature termination codons (PTCs), which may lead to mRNA degradation by the surveillance system known as nonsense-mediated mRNA decay (NMD) (Kurihara et al., 2009; Kalyna et al., 2012). Therefore, AS also regulates transcript abundance. Diverse environmental cues, including light and temperature, trigger rapid changes of splicing profiles (Wu et al., 2014; Calixto et al., 2018; Cheng and Tu, 2018). A growing number of factors involved in splice site definition, especially for U1 and U2 snRNP-related components, have been shown to play fundamental roles in relaying environmental signals to the splicing machinery. For example, light signals are transmitted to the U1 snRNP component Pre-mRNA-PROCESSING FACTOR 39-1 (PRP39) in the moss *Physcomitrium* (*Physcomitrella*) *patens* and to the U2 snRNP-associating factors SPLICING FACTOR FOR PHYTOCHROME SIGNALING (SFPS) and REDUCED RED LIGHT RESPONSES IN *cry1 cry2* BACKGROUND 1 (RRC1) in Arabidopsis (Shikata et al., 2012; Shikata et al., 2014; Xin et al., 2017; Shih et al., 2019; Xin et al., 2019; Lin et al., 2020), via the red/far-red light photoreceptor phytochromes (phys). AS also plays essential roles in temperature signaling, in particular for the regulation of flowering time in response to changes in temperature (Capovilla et al., 2015). At lower temperatures of 16°C, the transition from vegetative to reproductive development is blocked by the repressive complex composed of the two MADS-box transcription factors SHORT VEGETATIVE PHASE (SVP) and FLOWERING LOCUS M (FLM) (Lee et al., 2013; Posé et al., 2013). Increasing the temperature to 23°C triggers the degradation of SVP and reduce *FLM* transcript levels via AS regulation, coupled with NMD. Among *FLM* isoforms, *FLM-β* contributes the most to FLM function at low and ambient temperatures, as it can be translated into a functional FLM protein. Higher temperatures of 27 °C further promote the accumulation of *FLM-δ*, which is translated into a dominant-negative form of FLM, as well as numerous AS isoforms containing pre-mature termination codon (PTC) in the total pool of *FLM* transcripts, leading to relatively lower levels of functional FLM. While several U2 snRNP-related factors have been described as functioning during AS regulation of *FLM* (Lee et al., 2017; Park et al., 2019; Lee et al., 2020), the detailed biochemical and molecular mechanism still remains largely unclear.

As the earliest participant in pre-mRNA splicing, U1 snRNP is considered a focal point for AS regulation (Ule and Blencowe, 2019). U1 snRNP can be separated into U1 core and U1 auxiliary subunits, with the U1 core mediating the interaction between U1 snRNA and the 5’-SS and the U1 auxiliary subunits stabilizing this interaction. The plant U1 snRNP complex has yet to be purified biochemically, but homology searches have revealed possible plant counterparts to animal U1 core and several U1 auxiliary proteins (Wang and Brendel, 2004). The U1 auxiliary component Nuclear accommodation of mitochondria 8 (Nam8), comprising three RNA recognition motifs (RRMs), is involved in the regulation of meiotic pre-mRNA splicing, and is unnecessary for mitotic growth of yeast (*Saccharomyces cerevisiae*) (Spingola and Ares, 2000; Qiu et al., 2011). The human Nam8 homolog, T-cell intracellular antigen-1 (TIA-1), facilitates weak 5’-SS recognition by binding to U-rich sequences downstream of the 5’-SS and U1C (Förch et al., 2002; Wang et al., 2014; Li et al., 2017). TIA-1 is also an important factor mediating the formation of stress granules via its Gln-rich C-terminal region (Gilks et al., 2004). Mutations of TIA-1 in humans are associated with multiple pathologies, suggesting an essential regulatory role for Nam8/TIA1 in maintaining regular growth for eukaryotic cells (Hackman et al., 2013; Sánchez-Jiménez and Izquierdo, 2015). Although the function of Nam8/TIA1 in pre-mRNA splicing is well established in animals, little is known about the plant Nam8/TIA1 homolog. In Arabidopsis, members of the RNA Binding Protein 45/47 (RBP45/47) family, RBP45a-d and RBP47a-c’, showed high sequence similarity to Nam8/TIA-1 and have been proposed to function in stress granule formation and in hypoxia or ozone stress responses (Peal et al., 2011; Bhasin and Hülskamp, 2017; Kosmacz et al., 2019). However, whether these factors play roles in pre-mRNA splicing is unclear.

Although increasing evidence has indicated that regulation of pre-mRNA splicing is critical for proper flowering, the underlying components and the associated mechanism are little understood. In this study, we identified RBP45d as an evolutionarily conserved component of U1 snRNP. As the Arabidopsis ortholog of Nam8/TIA1, RBP45d associated with PRP39a to coordinately control the splicing of a specific set of genes. Plants lacking either RBP45d or PRP39a function flowered later than wild type. We also revealed that RBP45d and PRP39a regulate the temperature-sensitive AS of the floral repressor *FLM* through direct association with its pre-mRNA. Loss of proper *FLM* AS in the *rbp45d* and *prp39a* mutants resulted in the temperature insensitivity of flowering, and the mutants flowered as late at 28°C as they did at 22°C. Our study establishes RBP45d as an U1snRNP component in plants and provides new understanding of the role of U1 snRNP in ambient temperature-mediated flowering regulation by fine-tuning AS of *FLM*.

## RESULTS

### RBP45d is the Arabidopsis ortholog of yeast Nam8

Although they share high similarity with the splicing factor Nam8, little is known about the role of RBP45/47 family members on pre-mRNA splicing. To begin our investigation of the relationship between the Arabidopsis RBP45/47 family and Nam8, we aligned and compared their protein sequences (Supplemental Figure S1). The three RNA recognition motifs (RRMs) in Nam8 and RBP45/47 members were nearly identical, especially over the ribonucleoprotein (RNP) sub-domains RNP1 and RNP2. Outside of these conserved RRMs, we also noticed that Nam8 and RBP45d share similar sequence features, with a shorter N terminus and a longer C terminus compared to the other RBP45/47 members. The long Nam8 C-terminal region was demonstrated to act as a bridge for its interaction with PRP39 and U1C in yeast (Li et al., 2017; Plaschka et al., 2018). Based on this structural information, we speculated that the long C terminus might be required for RBP45d to associate with other components of the U1 snRNP in Arabidopsis. To test this hypothesis, we first examined the interaction between RBP45/47 family members and the Arabidopsis U1 snRNP component PRP39a, which was shown to regulate pre-mRNA splicing (Kanno et al., 2020), by yeast two-hybrid (Y2H) assay. RBP45d was the only member of the RBP45/47 family able to interact with PRP39a (Figure 1A). We further refined the region within RBP45d that mediates the interaction with PRP39a, which revealed that RBP45d forms a complex with PRP39a through its long C terminus, as expected (Supplemental Figure S2).

**Figure 1.**
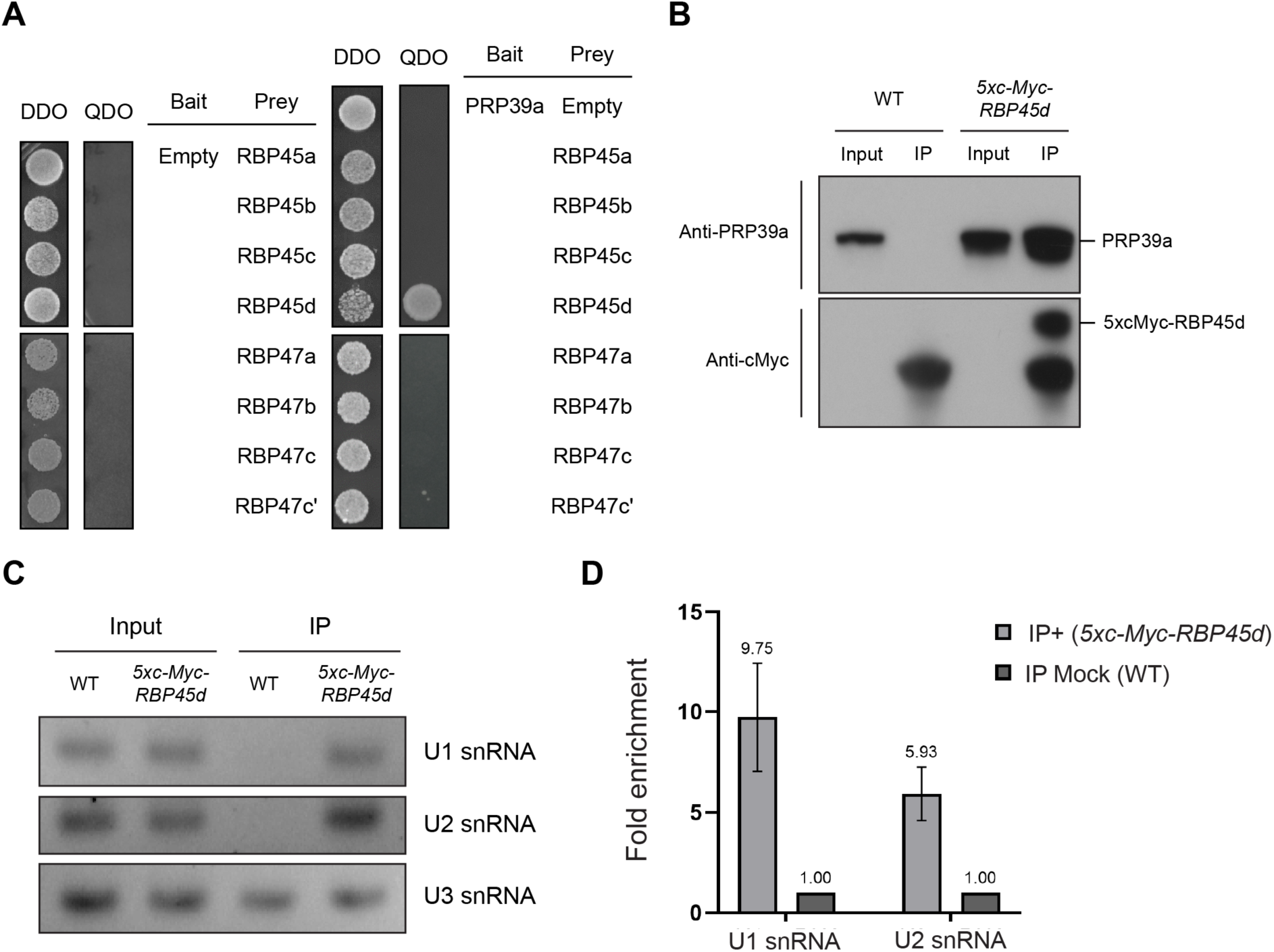
RBP45d is a plant U1 snRNP component. **(A)** RBP45d is the only RBP45/47 family member that interacts with PRP39a. For yeast two-hybrid (Y2H) assay, yeast cells co-transformed with constructs expressing *PRP39a* and *RBP45/47* family member proteins were diluted to an OD_600_ of 0.1 before plating onto synthetic defined (SD) medium lacking Leu and Trp (DDO, double dropout) for selection and SD medium lacking Leu, Trp, His and Ade (QDO, quadruple dropout) for interaction tests. Empty bait vector (pGBKT7; Empty) was co-transformed with prey clones as control. **(B)** Co-immunoprecipitation (co-IP) of RBP45d and PRP39a. Co-IP was performed using anti-c-Myc antibody to pull down associated protein complexes from lysates of Arabidopsis seedlings expressing *5x c-Myc-RBP45d*. Lysates from the wild-type (WT) Col-0 were used as negative control. Input corresponds to 50 µg of total lysate. Immunoprecipitated proteins were detected by immunoblotting with anti-cMyc and anti-PRP39a antibodies. **(C)** RNA immunoprecipitation (RIP) analysis of the association between RBP45d and snRNPs. Co-IP was performed using anti-c-Myc antibody and the immunoprecipitated complex was assessed for U1 snRNA and U2 snRNA abundance by end-point RT-PCR. WT lysates were used as controls. The non-specific associating transcript U3 snRNA was used as internal control. **(D)** Quantification of U1 snRNA and U2 snRNA by RT-qPCR, from the samples shown in (**C**).

The importance of this long C-terminal region prompted us to ask whether this feature of RBP45d was evolutionally conserved. Accordingly, we assembled protein sequences for 84 RBP45/47-like proteins from 19 representative species to perform a sequence alignment and phylogenetic analysis. All species had at least one RBP45/47 member with a long C terminus region (Supplemental Figure S3 and Supplemental Dataset S1). These C termini appeared highly conserved across species, suggesting their evolutionary importance (Supplemental Figure S3). We therefore defined these proteins as RBP45d-like.

Phylogenetic analysis indicated that RBP45/47-like proteins can be clearly separated into six major groups, which we named based on the closest Arabidopsis protein within each group. We also identified one additional group with eight Poaceae-specific RBP45/47-like members and lacking an Arabidopsis representative (Supplemental Figure S4A). Unlike RBP45d-like proteins, other RBP45/47 members had no clear putative orthologs in non-vascular plants (Supplemental Figure S4B), suggesting that *RBP45d*-like genes may have evolved earlier than other *RBP45/47* members. Taken together, although very similar in their protein sequence, RBP45/47 members can be divided into different functional groups. As the evolutionarily earliest RBP45/47 member and the most similar to Nam8, RBP45d may have inherited the role of Nam8 as an U1 snRNP component in Arabidopsis.

### RBP45d associates with the U1 snRNP in vivo

To further determine whether RBP45d is a component of U1 snRNP in Arabidopsis, we tested the interaction of RBP45d and PRP39a in vivo by co-immunoprecipitation (Co-IP) assays. Indeed, RBP45d effectively pulled down PRP39a from protein extracts of Arabidopsis seedlings, supporting the association between RBP45d and the U1 snRNP (Figure 1B). We also tested whether RBP45d interacted with other U1snRNP components in Arabidopsis by Y2H assay. As shown in Supplemental Figure S2A, RBP45d also associated with the U1 core components U1C and U1-70K, again via its C terminus. These findings are consistent with the hypothesis that RBP45d possesses the Nam8 functions of interacting with U1 components.

Another important feature of U1 snRNP components is their association with U1 snRNA (Terzi and Simpson, 2009; de Francisco Amorim et al., 2018). To test whether RBP45d came in proximity to U1 snRNA in vivo, we performed an RNA immunoprecipitation (RIP) assay with transgenic seedlings accumulating a c-Myc-tagged version of RBP45d in the Col-0 (WT) background. When compared to mock IP experiments performed on WT, the amount of immunoprecipitated U1 snRNA was enriched 10-fold in *5x c-Myc-RBP45d* transgenic seedlings relative to the amount of U3 snRNA, which serves as an internal control in IP experiments (Terzi and Simpson, 2009) (Figure 1C and 1D). Notably, we also detected a six-fold enrichment for the U2 snRNA, indicating that RBP45d is also in the vicinity of U2 snRNP. These findings agree with the latest observations of the yeast pre-spliceosome A complex (Plaschka et al., 2018) and the human pre-B complex cryogenic electron microscopy (cryoEM) structure (Charenton et al., 2019), which showed that U1 snRNP forms a transient complex with U2 snRNP (Plaschka et al., 2018; Charenton et al., 2019). We conclude here that RBP45d is a newly identified component of U1 snRNP in Arabidopsis.

### Flowering time is delayed in both *rbp45d* and *prp39a* mutants

To further investigate the role of *RBP45d* in Arabidopsis, we generated a null *rbp45d* mutant by CRISPR/Cas9-mediated genome editing. We identified T_1_ mutant plants harboring a 923-bp deletion between the two sgRNA target sites in the *RBP45d* coding region; we then collected T_2_ seeds and screened for T_2_ plants homozygous for the deletion and free of Cas9 (Supplemental Figure S5A-S5C). Coincidently, we also obtained two additional *rbp45d* mutants identified during a forward genetic screen in search for factors involved in pre-mRNA splicing (Kanno et al., 2020). Because these two *rbp45d* mutants were named *rbp45d-1* and *rbp45d-2*, we designated the genome-edited allele *rbp45d-3*. We confirmed the effects of the *RBP45d* deletion on *RBP45d* transcript levels by RT-PCR and RT-qPCR (Supplemental Figure S5D and S5E), indicating that *rbp45d-3* is likely a null allele.

The morphology of the *rbp45d-3* mutant was similar to that of wild type (WT) in both long day (LD) and short day (SD) conditions at 22°C, although the mutant flowered later than WT (Figure 2), similar to the phenotype of *prp39a* mutants (Wang et al., 2007; Kanno et al., 2017). The *rbp45d-1* and *rbp45d-2* mutants also exhibited delayed flowering relative to their WT when grown in LD and SD conditions (Supplemental Figure S6A and S6B). Importantly, the late flowering phenotype of *rbp45d-3* was complemented a transgene consisting of the *RBP45d* coding region under the control of the *RBP45d* promoter (Supplemental Figure S6C and S6D). Mutants with delayed flowering in both LD and SD conditions can be assigned to the autonomous pathway (Wu et al., 2020). Many of the factors in this pathway, including PRP39a, are involved in RNA processing and promote flowering by affecting transcript levels of the flowering repressor *FLOWERING LOCUS C* (*FLC*). This regulation is mainly mediated through the polyadenylation and splicing on a set of antisense long noncoding RNAs named *COOLAIR* produced from the *FLC* locus (Swiezewski et al., 2009; Marquardt et al., 2014). We therefore performed RT-qPCR to measure *FLC* and *COOLAIR* transcript levels in the *rbp45d-3* and *prp39a-1* mutants. However, even though we did detect higher *FLC* transcript levels in both *rbp45d-3* and *prp39a-1*, there was no significant difference for *COOLAIR* transcript levels (Supplemental Figure S7A, S7B) (Marquardt et al., 2014). This result indicates that RBP45d and PRP39a might promote flowering through other unknown pathways.

**Figure 2.**
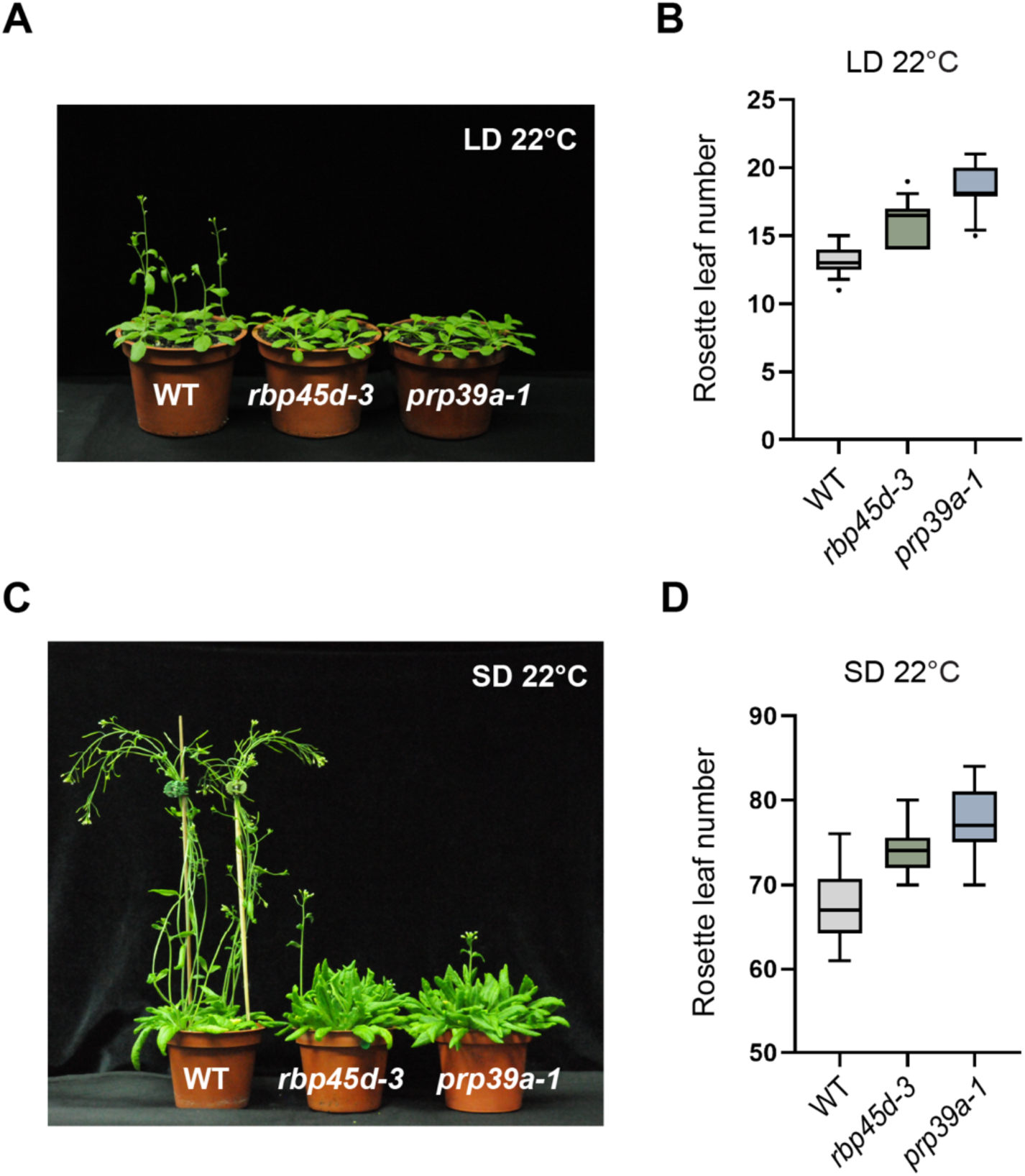
RBP45d and PRP39a promote flowering independently of the photoperiod. **(A)** Representative photographs of WT (Col-0), *rbp45d-3* and *prp39a-1* plants grown in long day (LD) conditions. **(B)** Mean rosette leaf number of WT (Col-0), *rbp45d-3* and *prp39a-1* plants at bolting in LD conditions. **(C)** Representative photographs of WT, *rbp45d-3* and *prp39a-1* plants grown in short day (SD) conditions. **(D)** Mean rosette leaf number of WT (Col-0), *rbp45d-3* and *prp39a-1* plants at bolting in SD conditions. At least 12 plants were scored and the experiments were performed for at least three times.

### RBP45d and PRP39a together regulate pre-mRNA splicing of a subset of genes

To gain a better understanding of how RBP45d and PRP39a, as part of the U1 auxiliary subunits, regulate splicing and flowering, we performed a transcriptome deep sequencing (RNA-seq) to analyze the expression and splicing profiles of all Arabidopsis genes in the *rbp45d-3* and *prp39a-1* mutant backgrounds. A comparison with the WT transcriptome at the same developmental stage highlighted only 97 and 121 differentially expressed genes (DEGs) in *rbp45d-3* and *prp39a-1*, respectively (Figure 3A), suggesting that the loss of RBP45d or PRP39a has no widespread influence on transcriptional regulation. By contrast, we identified abundant differential AS (DAS) events in both mutants (Figure 3A), supporting a main role for these two factors in pre-mRNA splicing. Interestingly, we noticed far more differential intron retention (DIR) and exon skipping (DES) events in both mutants compared to WT (Figure 3B). This result indicated that RBP45d and PRP39a may facilitate the recognition of cryptic splice sites (SSs), and thus the loss of either factor leads to a weaker recognition of cryptic SSs, eventually resulting in IR or ES.

**Figure 3.**
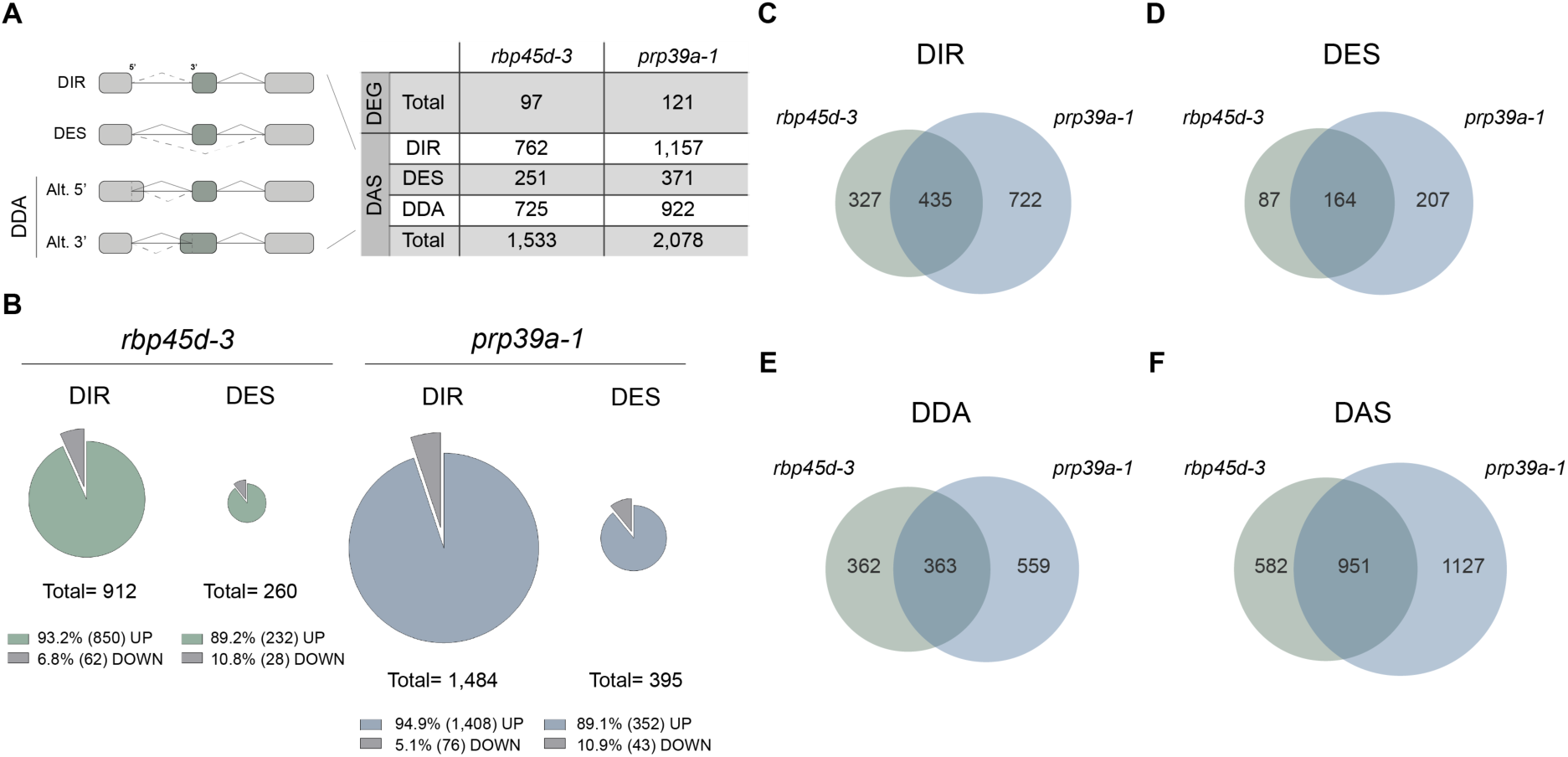
RBP45d and PRP39a co-regulate pre-mRNA splicing for a specific set of genes. (**A**) Numbers of differentially expressed genes (DEGs) and differential alternative splicing (DAS) events, including differential intron retention (DIR), differential exon skipping (DES) and differential alternative 5’- or 3’-splice site (DDA) identified in the *rbp45d-3* and *prp39a-1* mutants relative to the WT Col-0. **(B)** Pie charts showing the percentage of up- and downregulated DIR and DES events in each mutant. (**C**-**F)** Venn diagram showing the number of unique or shared DIR, DES, DDA events and total DAS genes to *rbp45d-3* and *prp39a-1* mutants.

Based on the observations that *RBP45d* and *PRP39a* expression profiles are highly correlated (Supplemental Figure S8), RBP45d and PRP39a form a complex, and that loss of either factor results in similar phenotypes, we suspected that the two factors may co-regulate AS. Indeed, over 57% (435 of 762) of genes exhibiting DIR in the *rbp45d-3* mutant also behave similarly in the *prp39a-1* mutant (Figure 3C). We obtained similar overlaps for DES and differential alternative 5’- or 3’-SSs (DDA) events, with 65% (164 of 251) of DES and 50% (363 of 725) of DDA genes in *rbp45d-3* mutant also affected in the *prp39a-1* mutant (Figure 3D and 3E). Overall, 62% (951 of 1,533) of DAS genes in the *rbp45d-3* mutant were also misregulated in the *prp39a-1* mutant (Figure 3F). These results strongly suggest that the two splicing factors function together in fine-tuning pre-mRNA splicing for the same group of genes.

To better understand the characteristic of DAS genes detected in both mutants, we conducted a gene ontology (GO) and UniProt-keyword (KW) enrichment analysis on gene sets co-regulated by RBP45d and PRP39a. GO analysis revealed that the most significantly enriched term in both gene sets is “chloroplast” (GO: 0009507) (Supplemental Figure S9A and S9B). In line with this result, three related keywords (transit peptide, chloroplast, and plastid) were highly enriched in the UniProt keyword analysis (Supplemental Dataset S2), indicating that the RBP45d and PRP39a complex regulates AS for genes encoding proteins targeted to the chloroplast.

We next examined the property of DIR genes in both mutants, including their length and the number of introns in genes affected by IR. These differentially retained introns were generally shorter in length compared to that of retained introns that do not depend on RBP45d or PRP39a (Supplemental Figure S9C and S9D). From the analysis of intron number, we discovered that DIR in *rbp45d-3* and *prp39a-1* mutants tends to occur for genes with more introns compared to IR genes that are not dependent on RBP45d or PRP39a (Supplemental Figure S9E and S9F). In summary, RNA-seq analysis, revealed that RBP45d and PRP39a co-regulate pre-mRNA splicing for a subset of introns that tend to be shorter in length and located in genes with high numbers of introns.

### *RBP45d* regulates AS of *PRP39a*

The genes encoding splicing regulators often undergo intensive AS regulation themselves (Verhage et al., 2017). In the current Araport 11 annotation, *PRP39a* is represented by two major AS isoforms that differ in the incorporation of exons 6 and 7 (Figure 4A). However, our RNA-seq analysis revealed a more complicated AS pattern for *PRP39a*. We noticed substantial AS involving intron 4 and regions between exon 6 and 7. Interestingly, these AS isoforms were not detected in *rbp45d-3* (Figure 4A). Based on RT-PCR and Sanger sequencing of PCR products, we identified at least seven distinct *PRP39a* transcript isoforms (Figure 4B). We also observed aberrant AS regulation of *PRP39a* transcripts in the *rbp45d-1* and *rbp45d-2* mutants (Supplemental Figure S10A) and in the *prp39a* mutant in a previous study (Kanno et al., 2017). Together, we conclude that the proper splicing of *PRP39a* requires both U1 auxiliary factors RBP45d and PRP39a.

**Figure 4.**
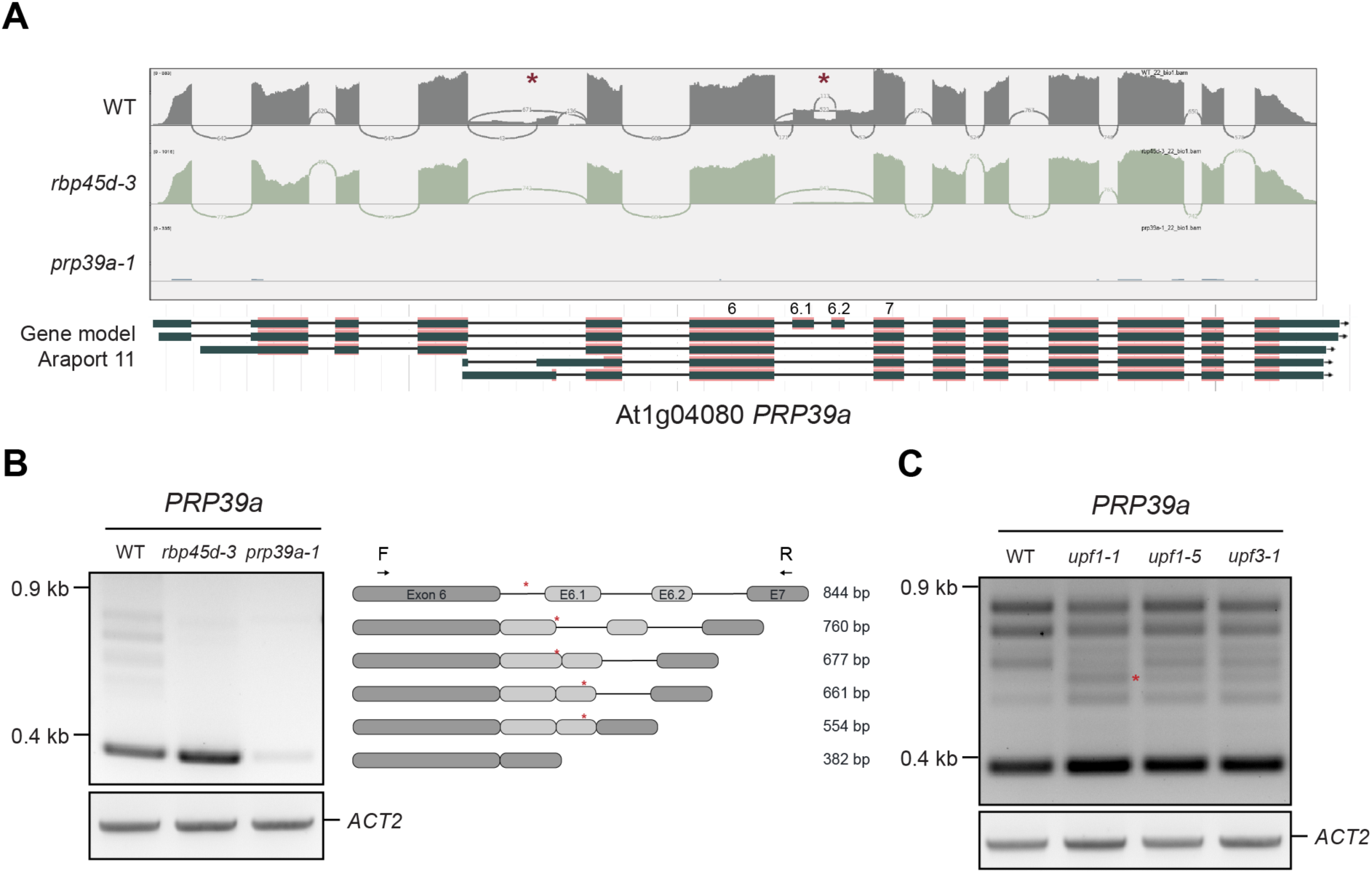
RBP45d regulates *PRP39a* pre-mRNA splicing. (**A)** Coverage plot showing the reads mapping to the *PRP39a* locus in WT (Col-0), *rbp45d-3* and *prp39a-1.* The plots were generated for one representative out of the three biological replicates for RNA-seq. Sashimi plots were generated in IGV. Red asterisks indicate the regions with significant differential AS between WT and *rbp45d-3*. Gene models for *PRP39a* were from the Araport 11 annotation. **(B)** End-point RT-PCR analysis of *PRP39a* AS in WT, *rbp45d-3* and *prp39a-1* (left). *ACTIN 2* (*ACT2*) was used as internal control. A schematic representation of the corresponding bands amplified by RT-PCR are shown on the right, with primer positions used for PCR and pre-mature termination codon resulting from AS (red asterisks) indicated. **(C)** End-point RT-PCR analysis of *PRP39a* AS in WT and the NMD mutants *upf1-1*, *upf1-5* and *upf3-1*. The red asterisk indicates the band enriched in *upf1-1*, *upf1-5* and *upf3-1* compared to WT. *ACT2* was used as internal control.

AS of *PRP39* pre-mRNA was also recently observed in mammalian systems (De Bortoli et al., 2019). In the case of murine (*Mus musculus*) and human *PRP39*, the inclusion of an alternative exon between exons 8 and 9 produces a PTC-containing isoform, which further leads to its degradation by NMD or production of a non-functional, truncated PRP39 protein that lacks the C terminus and thus disrupts its function as a backbone of U1 auxiliary complex. We determined that the Arabidopsis *PRP39a* pre-mRNA undergoes more complex AS regulation than its mammalian counterpart. Notably, all detected isoforms contained PTCs in the middle of the transcript, as in the mammalian system. We observed the accumulation of several isoforms in the NMD mutants *upf1-1*, *upf1-5* and *upf3-1* (Figure 4C), as might be expected for isoforms targeted by the NMD pathway. We obtained similar results with seedlings treated with cycloheximide (CHX), a translation inhibitor that blocks NMD (Carter et al., 1995), again supporting the targeting of some isoforms to the NMD pathway (Supplemental Figure S10B). Taken together, both RBP45d and PRP39a itself participate in the AS regulation of *PRP39a* pre-mRNA, possibly serving as an auto-regulatory mechanism for the splicing machinery.

### RBP45d regulates pre-mRNA splicing by direct association with its RNA targets

To understand how RBP45d regulates AS and to identify its potential targets, we performed RNA immunoprecipitation (RIP) and sequenced the RNAs associated with RBP45d. RIP sequencing (RIP-seq) analysis identified 4,438 transcripts enriched in at least two of the three biological repeats as RBP45d targets (Figure 5A; Supplemental Dataset S3). A comparison between the lists of 4,438 RBP45d target genes from RIP-seq and 97 DEGs from RNA-seq data yielded a very limited overlap of three genes (Supplemental Figure S11), supporting the notion that DEGs in *rbp45d-3* are not direct targets of RBP45d. By contrast, more than 42% (646 of 1,533) of DAS genes were also identified as RBP45d target genes by RIP-seq (Figure 5B). This finding suggested that many of the DAS cases in *rbp45d-3* are regulated directly by RBP45d. In the case of *PRP39a*, we also detected enriched AS peaks in its coding region for two of the three RIP-seq biological replicates (Figure 5C). The interaction between RBP45d and *PRP39a* transcripts was further validated by RT-qPCR (Figure 5D), confirming that *PRP39a* transcripts are in vivo targets of RBP45d.

**Figure 5.**
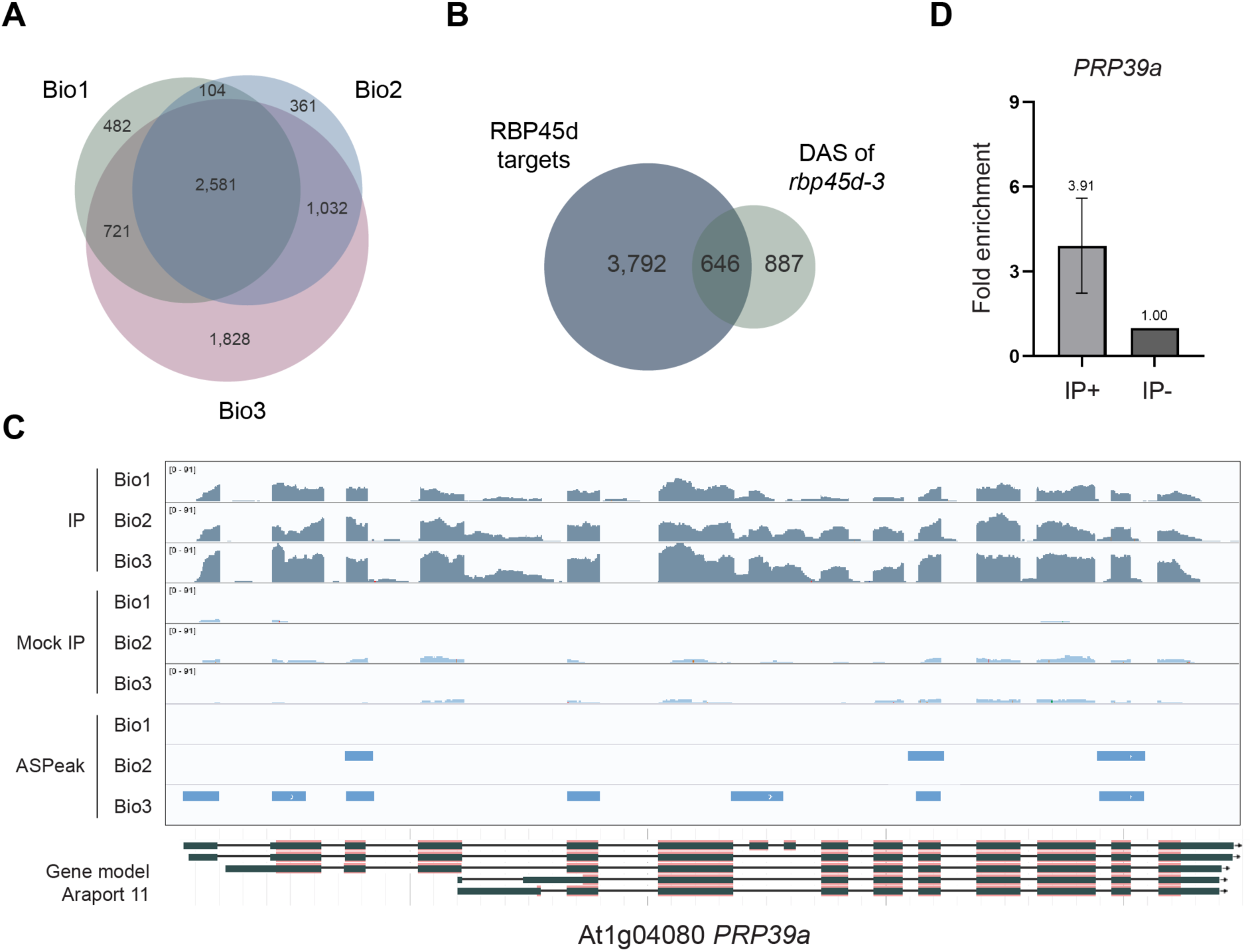
RBP45d regulates pre-mRNA splicing by direct association with its RNA targets. **(A)** Venn diagram showing the overlap between genes enriched across the three biological RIP-seq replicates. Transcripts enriched in at least two of the three biological replicates were considered as RBP45d targets. **(B)** Venn diagram showing the extent of overlap between RBP45d targets identified by RIP-seq and DAS genes in the *rbp45d-3* mutant. (**C)** Coverage plot showing RIP-seq reads mapping to *PRP39a*, as visualized with IGV for each of the three biological replicates from immunoprecipitated RNA (IP) and Mock IP. Light blue rectangles in the lower three lanes indicate peak enrichment, as determined by ASPeak software. The *PRP39a* gene model is from the Araport 11 annotation. **(D)** RT-qPCR validation of the association between RBP45d and the *PRP39a* transcript. The numbers above the bars indicate the fold-enrichment using U3 snRNA as internal control. Error bars represent standard error of the mean (*n* = 4).

### Temperature-responsive AS of *FLM* is regulated by RBP45d and PRP39a

As described above, RBP45d and PRP39a are involved in the autonomous flowering pathway, as their late flowering phenotype is photoperiod-independent and they accumulate higher *FLC* transcripts (Supplemental Figure S7A and S12A) (Wang et al., 2007; Wu et al., 2020). However, the transcript levels of other autonomous pathway genes remained unaffected (Supplemental Figure S12B), suggesting that the two splicing factors may influence flowering through other mechanisms. Intriguingly, we observed higher levels and an aberrant splicing pattern for the flowering repressor *FLOWERING LOCUS M* (*FLM*) in both mutants compared to the WT (Figure 6A; Supplemental Figure S12C). *FLM* has effects comparable to those of *FLC* in floral repression (Ratcliffe et al., 2001; Scortecci et al., 2001), and plays key roles specifically in temperature-dependent regulation of flowering, as changes in ambient temperature strongly affect the level and splicing pattern of *FLM* transcripts (Balasubramanian et al., 2006). In both *rbp45d-3* and *prp39a-1* mutants, we detected higher levels of isoform *FLM-β*, which encodes a functional repressor, compared to WT, while the levels of *FLM-δ*, encoding a dominant-negative version of FLM, were reduced (Figure 6B). Thus, when grown at 22°C, *FLM-β* in these mutants represented a much higher fraction of bulk *FLM* transcripts compared to the WT, in agreement with the late flowering phenotype of these two mutants. In addition, RT-PCR analysis revealed aberrant *FLM* splicing patterns in both mutants (Figure 6C), suggesting that RBP45d and PRP39a regulate AS of *FLM* transcripts. This regulation was likely mediated by direct physical interaction between RBP45d and *FLM* transcripts in vivo, as determined by RIP-seq (Figure 6A) and further validated by RIP and RT-qPCR (Figure 6D). These results strongly support a role for RBP45d in modulating flowering time through its association with *FLM* transcripts to regulate their splicing preference.

**Figure 6.**
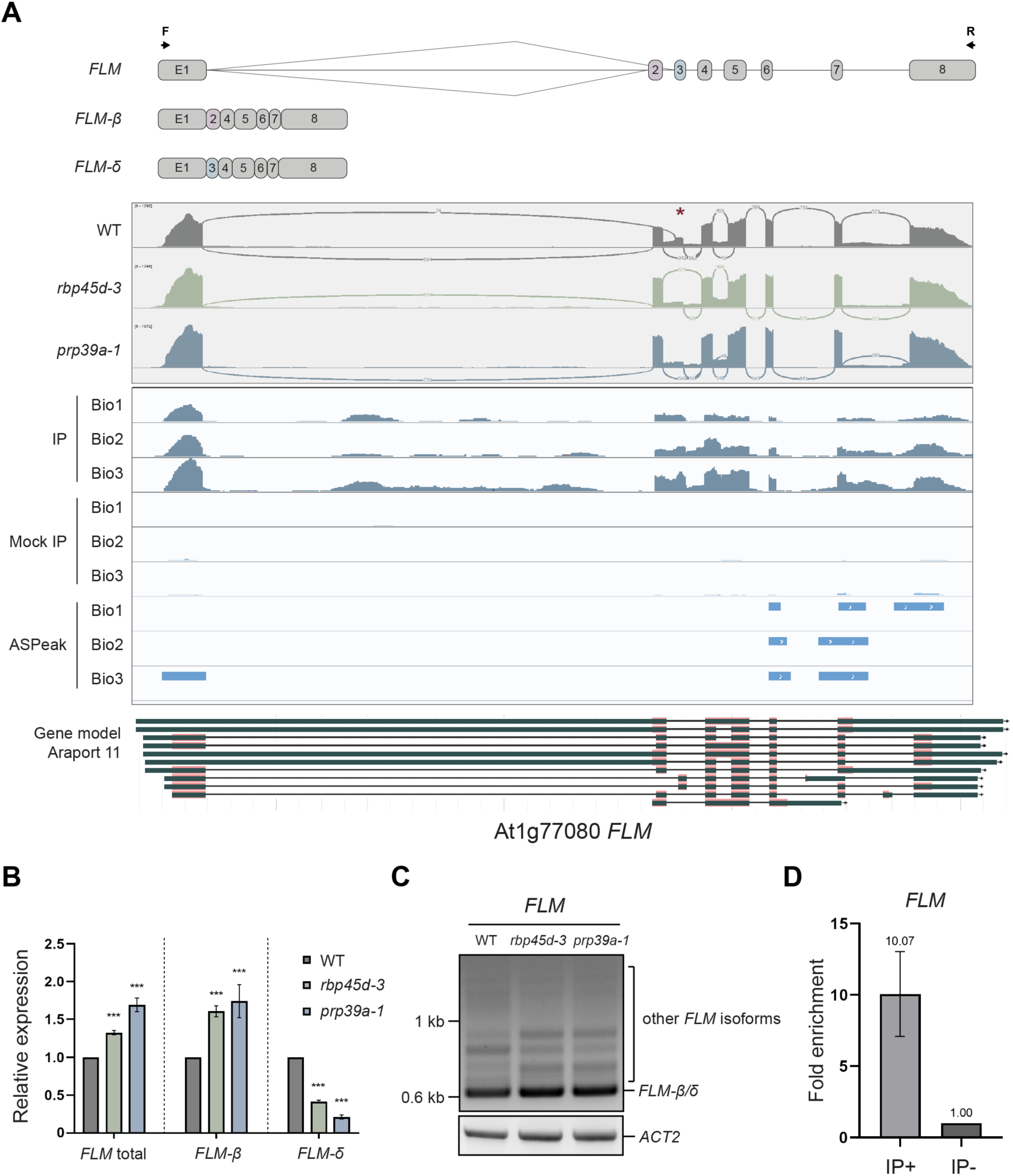
RBP45d and PRP39a contribute to AS regulation of the flowering repressor *FLM*. **(A)** Alternative splicing (AS) at FLM in WT (Col-0), *rbp45d-3* and *prp39a-1*. Top: schematic representation of *FLM* splice variants. Exons present only in the *FLM-β* and *FLM-δ* isoforms are labeled. Coverage plots (middle plots with gray background) show reads mapping to *FLM*, visualized in IGV from one of the three RNA-seq biological replicates for WT, *rbp45d-3* and *prp39a-1*. Sashimi plots were generated in IGV. The red asterisk indicates the region of significant differential AS between WT and *rbp45d-3* or *prp39a-1*. Bottom panel shows coverage plots (in IGV) of RIP-seq reads mapping to *FLM.* Light blue rectangles in the lower three lanes indicate enriched peaks, as determined by ASPeak. **(B)** RT-qPCR of bulk *FLM* transcript levels and for its two major isoforms *FLM-β* and *FLM-δ* in WT, *rbp45d-3* and *prp39a-1*. Error bars represent standard error of the mean (*n* = 3). ** *P* < 0.01; *** *P* < 0.001, as determined by Student’s *t*-test. *PP2AA3* was used as internal control. **(C)** End-point RT-PCR analysis of *FLM* AS in WT, *rbp45d-3* and *prp39a-1*. *ACTIN 2* (*ACT2*) was used as internal control. The primers (F and R) used are indicated on the *FLM* gene structure in panel (**A**). **(D)** RT-qPCR validation of the association between RBP45d and *FLM* transcripts. The numbers above the bar indicate the fold-enrichment using U3 snRNA as internal control. Error bars represent standard error of the mean (*n* = 3).

The transcript levels of *FLM-β* decrease as temperatures rise, concomitantly with an increase in *FLM-δ*, which raises the ratio of non-functional *FLM* isoforms in the pool of *FLM* transcripts (Sureshkumar et al., 2016). Since RBP45d and PRP39a are involve in maintaining proper splicing of *FLM* (Figure 6C), both factors may also affect the temperature-responsive AS of *FLM*. We therefore examined the levels of *FLM* isoforms by RT-PCR in Arabidopsis plants always grown at 22°C or shifted from 22°C to 28°C for 24 h. In agreement with our hypothesis, *FLM* transcript patterns in both *rbp45d-3* and *prp39a-1* mutants shifted from 22°C to 28°C were distinct from those of the WT (Figure 7A). In both mutants, the higher temperature resulted in a drop in *FLM-β/δ* levels, as evidenced by RT-PCR, that was not as pronounced as that seen in the WT. The generation of other non-functional isoforms was also much reduced in both mutants. We observed a similar phenomenon when shifting plants from 16°C to 28°C (Supplemental Figure S13). We thus concluded that RBP45d and PRP39a facilitate temperature-induced AS of *FLM*.

**Figure 7.**
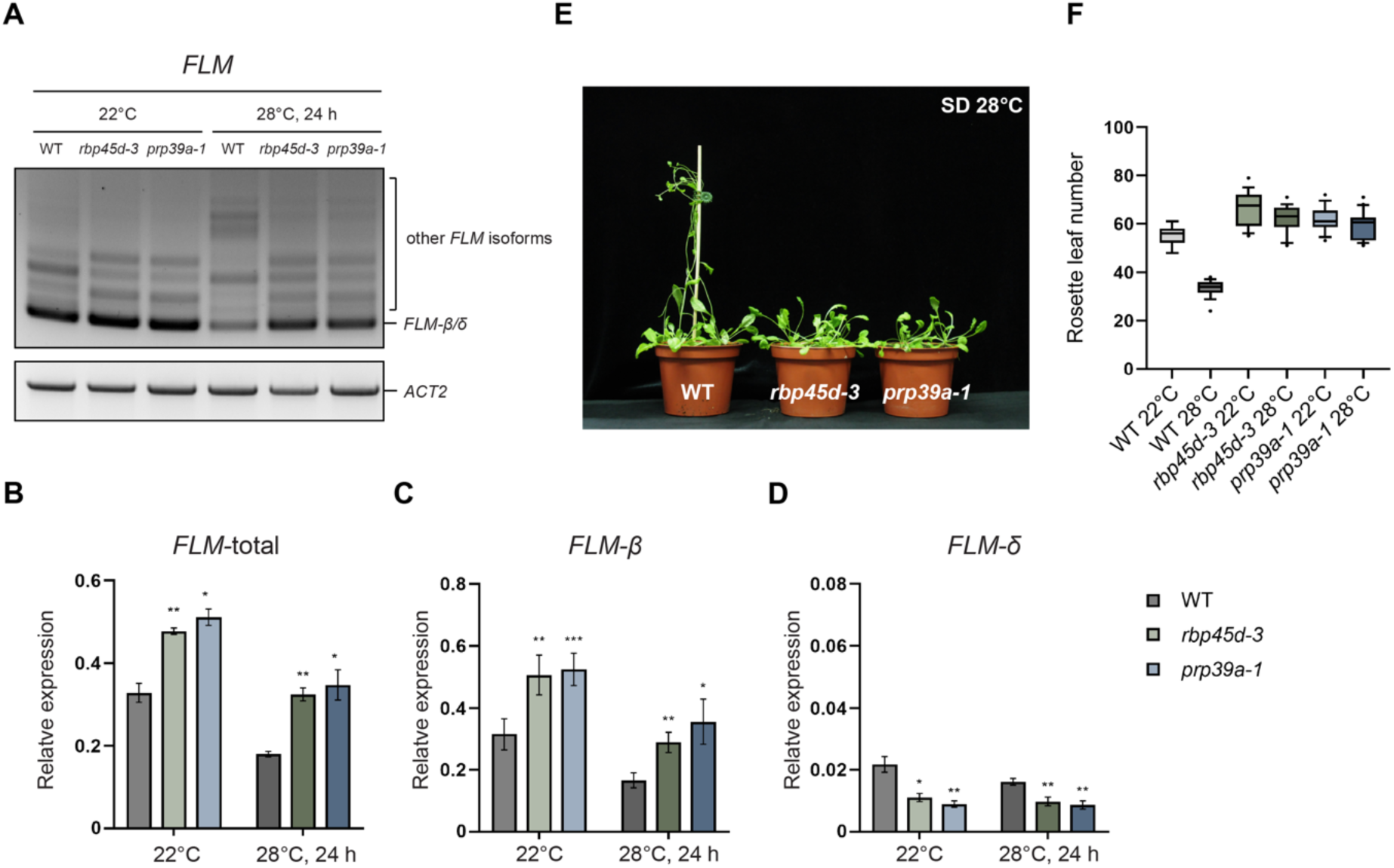
Temperature-responsive *FLM* AS is regulated by RBP45d and PRP39a. (**A)** End-point RT-PCR analysis of AS for *FLM* isoforms in WT, *rbp45d-3* and *prp39a-1* grown at 22°C or shifted from 22°C to 28°C for 24 h. *ACT2* was used as internal control. **(B-D)** RT-qPCR of bulk *FLM* transcript levels and for its two major isoforms *FLM-β* and *FLM-δ* in WT, *rbp45d-3* and *prp39a-1* grown at 22°C or shifted from 22°C to 28°C for 24 h. Error bars represent standard error of the mean. * *P* < 0.05; ** *P* < 0.01; *** *P* < 0.001, as determined by Student’s *t*-test. *PP2AA3* was used as internal control. **(E)** representative photographs of WT (Col-0), *rbp45d-3* and *prp39a-1* plants grown in SD conditions and shifted from 22°C to 28°C. **(F)** Mean rosette leaf number of WT (Col-0), *rbp45d-3* and *prp39a-1* plants at bolting when grown at 22°C or 28°C in SD conditions. At least 12 plants were scored and the experiments were performed for at three times.

We then analyzed the levels of *FLM-β* and *FLM-δ* isoforms individually by RT-qPCR. Although bulk *FLM* and *FLM-β* transcripts decreased at higher temperature, both bulk *FLM* and *FLM-β* isoforms accumulated to higher levels in the mutants compared to WT (Figure 7B and 7C). By contrast, *FLM-δ* transcripts remained at a low level that does not respond to the shift in temperature (Figure 7D). Together, these findings strongly suggest that flowering induced by higher temperatures may be abnormal in both mutants. Indeed, both *rbp45d-3* and *prp39a-1* mutants flowered later than WT when grown in SD conditions at 28°C (Figure 7E and 7F), with a near complete loss of thermal responsiveness, presumably caused by defects in regulating AS of *FLM* under changing temperature conditions.

## DISCUSSION

Alternative splicing (AS) is an essential regulatory step in many signaling pathways. However, the components involved and the underlying mechanism remain largely unknown in plants. Previous studies in yeast and mammalian systems have addressed the importance of the U1 snRNP component Nam8/Tia1-like proteins in pre-mRNA splicing, but their plant orthologs have yet to be described. In this study, we identified and characterized Arabidopsis RBP45d as a novel U1 snRNP component. From RNA-seq data, we identified a specific subset of AS events, including those for *FLM* and *PRP39a*, as being co-regulated by RBP45d and PRP39a. RIP experiments showed that RBP45d regulates AS events for *FLM* and *PRP39a* through direct association with their pre-mRNAs. Further investigation revealed that the aberrant pre-mRNA splicing pattern of *FLM* in *rbp45d* and *prp39a* mutants leads to their temperature-insensitive late flowering phenotype. We conclude that RBP45d is a component of the U1 snRNP and is critical for flowering time control in plants.

### RBP45d is a long neglected component of U1 snRNP

Despite their high similarity to the yeast U1 snRNP component Nam8, plant RBP45/47 family members have long been considered as factors involved in the formation of stress granules and stress responses (Peal et al., 2011; Bhasin and Hülskamp, 2017; Kosmacz et al., 2019) Among these proteins, RBP47b was even used as a marker of stress granules (Kosmacz et al., 2019) Interestingly, RBP45d was the only Nam8/TIA1 homolog in Arabidopsis that was neither detected in stress granules nor responsive to stress conditions, suggesting that RBP45d may have functions distinct from those of other RBP45/47 members. A protein sequence alignment between RBP45d and other RBP45/47 members highlighted that RBP45d lacks the glutamine (Gln)-rich N terminus and possesses a long C terminus with low Gln content conserved in other members, making it the closest Nam8 ortholog in Arabidopsis (Supplemental Figure S1, Supplemental Table S1A and S1B). Previous studies have shown that Gln-rich regions, or more precisely regions rich in Gln, Asp, Tyr, and Gly, form prion-related domains (PrD) with a high propensity to aggregate, and are thus important in nucleating mRNA-protein complexes during stress granule formation (Michelitsch and Weissman, 2000; Gilks et al., 2004; Kosmacz et al., 2019). The lack of this Gln-rich region may explain why RBP45d was not detected a component of stress granules in previous studies. Indeed, all RBP45/47 family members except Nam8 and RBP45d possessed PrD-like regions at either or both their N and C termini (Supplemental Figure S14), as predicted by the PrD prediction software PLAAC (Lancaster et al., 2014).

We showed the importance of the RBP45d C terminus in mediating the interaction with U1-snRNP. Nam8 also associates with the U1 auxiliary protein PRP39 through its long C terminus in yeast (Li et al., 2017). Both observations strongly support RBP45d function as an U1 component rather than as a stress granule nucleating factor. At this stage, we cannot exclude that other RBP45/47 family members also function in pre-mRNA splicing. However, when NpRBP45 and NpRBP47 were transiently overexpressed in *Nicotiana plumbaginifolia* protoplasts, no obvious differences in splicing of the reporter RNAs were observed (Lorković et al., 2000).

Phylogenetic analysis of RBP45/47 family members from representative species revealed that all species contain at least one ortholog harboring a short N terminus and a highly conserved long C terminus, which we named here RBP45d-like proteins (Supplemental Figure S3). Of note, while *RBP45d*-*like* genes are widely distributed among green plants, orthologs of other *RBP45/47* genes are missing in ancient land plants, and only emerged with eudicots (Supplemental Figure S4). We also discovered that *RBP45d-like* genes have undergone gene duplication, loss of the sequence encoding the C terminus, and gain of the sequence encoding the Gln-rich N terminus (Supplemental Dataset S1) over the course of evolution. We speculate that duplication may have presented an opportunity for proteins encoded by these *RBP45d-like* genes to lose the ability to associate with U1 snRNP, gaining polymerization ability, and gradually developing into stress granules nucleating factors. This observation offers the interesting possibility that other *RBP45/47* genes evolved from the U1 snRNP component RBP45d.

### RBP45d and PRP39a as novel factors involved in thermal responsive flowering regulation

Many factors such as SF1, U2AF65a, GLYCINE-RICH RNA-BINDING PROTEIN 7 (GRP7), GRP8, ARGININE-RICH CYCLIN 1 (CYCL1) and CYCLIN-DEPENDENT KINASE G2 (CDKG2), regulate AS of *FLM* (Lee et al., 2017; Park et al., 2019; Steffen et al., 2019; Lee et al., 2020; Nibau et al., 2020) by either facilitating or suppressing the accumulation of the functional *FLM-β* isoform. In our study, RBP45d and PRP39a suppressed the formation of *FLM-β*. Lack of either protein resulted in an accumulation of *FLM-β* that correlated with the late flowering phenotype. RIP experiments identified *FLM* as an in vivo target of RBP45d, suggesting that RBP45d and PRP39a are directly involved in the AS regulation of *FLM*. These findings pointed out that U1 snRNP components also participate in the regulation of *FLM* splicing. Rising temperatures quickly lowered the levels of the *FLM-β* isoform by modulating the splicing machinery, as evidenced by the generation of numerous non-functional PTC-containing isoforms. (Sureshkumar et al., 2016; Capovilla et al., 2017) (Figure 7A). The effect of rising temperatures on *FLM* AS was much milder in the *rbp45d* and *prp39a* mutants, suggesting that the RBP45d-PRP39a complex may mediate the effect of temperature on the splicing machinery. However, how temperature signals affect the activity of these splicing factors remains unknown.

### Convergent evolution of *PRP39* AS and its role in providing flexibility to splice site definition

During intron splicing, dynamic interactions not only occur between snRNPs but also within the complex. The degree of stability and flexibility exhibited by the complex may vary in different organisms. In the case of U1 snRNP, yeast U1 auxiliary proteins associate with the U1 core in a relatively stable interaction. However, in mammals, U1 auxiliary factors only transiently interact with the core complex (Li et al., 2017; De Bortoli et al., 2019). In plants, biochemical and structural information on the U1 core and auxiliary components is limited; it is therefore unclear whether U1 auxiliary factors, including PRP39a and RBP45d, stably or transiently associate with the U1 core. However, based on our observation that defects in U1 snRNP auxiliary components merely affect splicing of a few introns (Figure 3A), we propose that plant U1 auxiliary components, at least PRP39a and RBP45d, may function similarly to splicing regulators, associating dynamically with the U1 core and playing important roles in facilitating the recognition of weak or cryptic SSs within specific pre-mRNAs.

PRP39 forms homodimers in mammals, and PRP39 forms heterodimers with PRP42 in yeast, to bridge the interaction between U1 auxiliary and U1 core components (Li et al., 2017; Plaschka et al., 2018). *PRP39* pre-mRNA undergoes tissue-specific AS that results in truncated PRP39 proteins that can no longer dimerize (De Bortoli et al., 2019). In this study, we discovered that, like mammalian *PRP39*, AS of Arabidopsis *PRP39a* generates isoforms carrying PTCs in the middle of its transcripts. That such PTC-containing *PRP39* isoforms were detectable in both plant and mammalian systems suggests that *PRP39* is not a target of the NMD pathway. In agreement with this speculation, the effect of CHX in both systems was moderate. It is possible that PTC-containing *PRP39* isoforms might be translated into truncated PRP39a proteins that might then disrupt the normal function of U1 snRNP. The existence of both full-length and truncated PRP39 may provide some flexibility to the splicing machinery during SS selection. Although distant in evolution, both plant and animal systems developed similar regulatory mechanisms targeting the same gene independently, indicating this flexibility may be important and may be considered an example of molecular convergent evolution.

## MATERIAL AND METHODS

### Plant materials and growth conditions

All *Arabidopsis thaliana* genotypes used in this study were in the Columbia-0 (Col-0) background. The T-DNA insertion mutants *prp39a-1* (SALK_133733), *upf1-1* (*lba1*, CS6940) (Yoine et al., 2006), *upf1-5* (SALK_112922) and *upf3-1* (SALK_025175) were obtained from the Arabidopsis Biological Resource Center (ABRC). *rbp45d-3* was generated by genome editing via CRISPR-Cas9. Two sgRNA sequences were designed to target exon 1 or exon 2 of *RBP45d* with the web-based sgRNA design tool CRISPR-P v2.0 (http://crispr.hzau.edu.cn/CRISPR2/) (Liu et al., 2017). Transgenic plants were genotyped by PCR with specific primer pairs (Supplemental Data Set S4) to screen for the complete deletion between mutation sites targeted by two sgRNAs. Genomic DNA was extracted in hexadecyl trimethyl ammonium bromide (CTAB) buffer (100 mM Tris-HCl pH 8.0, 20 mM ethylenediaminetetraacetic acid [EDTA], 1.4 M NaCl, 1% [w/v] polyvinyl pyrrolidone [PVP] and 2% [w/v] CTAB). Homozygous Cas9-free T_3_ progeny were validated by RT-PCR, RT-qPCR and antibiotic selection on half-strength Murashige and Skoog (MS) medium containing 20 µg/mL hygromycin B Gold. *RBP45d-5xc-Myc* overexpression lines were generated by Agrobacterium (*Agrobacterium tumefaciens*)-mediated transformation using Agrobacterium strain GV3101 carrying the construct pCAMBIA1390_p35S::*RBP45d*-5xc-Myc. *rbp45d-1*, *rbp45d-2*, *prp39a-3* and *prp39a-4* were previously reported (Kanno et al., 2017; Kanno et al., 2020) For the generation of *rbp45d-3 RBP45dpro:RBP45d* lines, a 1.6-kb sequence upstream of the *RBP45d* transcription start site was cloned upstream of the *RBP45d* coding sequence in pCAMBIA1390; the resulting construct was introduced into the *rbp45d-3* mutant by Agrobacterium-mediated transformation.

### Phenotypic analysis

For flowering time, plants were grown on soil at 22°C in LD conditions (16-h light/8-h dark; light intensity 120 µmol m^−2^s^−1^), or at 22°C in SD conditions (8-h light/16-h dark; light intensity 120 µmol m^−2^s^−1^). Flowering time was recorded by measuring the number of rosette leaves when the inflorescence reached 1 cm. in height For flowering experiments at high temperatures, seedlings were first grown on soil at 22°C in LD conditions for 2 d, shifted to 22°C in SD conditions for 10 d and either maintained in SD conditions at 22°C or shifted to 28°C in SD conditions. For temperature treatment assays, seedlings were grown at either 16°C or 22°C in LD conditions for 12 d before a shift to 28°C in LD conditions at ZT13 (Zeitgeber 13, with lights on being ZT 0; corresponds to 3 h before lights off). Plant tissues were collected 24 h after temperature treatment for RNA extraction. For cycloheximide (CHX) treatment, Col-0 seedlings were grown in half-strength liquid MS medium for 7 d before being treated with either DMSO (solvent) or CHX (10 μg/mL) for 4 h.

### Yeast two-hybrid (Y2H) assays

Y2H experiments were performed using the Matchmaker^TM^ Gold Yeast Two-Hybrid System (Clontech). The coding sequences of *PRP39a* or *RBP45d* were cloned into the pGBKT7 vector in-frame with the coding sequence of the GAL4 DNA binding domain, while the coding sequences of *RBP45a-d*, *RBP47a-c’*, *U1A*, *U1C*, *U1-70K*, and various truncations of *RBP45d* were cloned into the pGADT7 vector in-frame with the coding sequence of the GAL4 activation domain. The resulting constructs were transformed into yeast strain Y2HGold (for pGBKT7 clones) or Y187 (for pGADT7 clones). Transformants were then mated on YPDA medium overnight and selected on synthetic defined medium lacking leucine and tryptophan (DDO; double dropout). Mated cells were incubated in DDO medium overnight before being diluted by the indicated factors before being spotted onto both DDO medium and synthetic defined medium lacking leucine, tryptophan, histidine and adenosine (QDO; quadruple dropout). Yeast cells were grown 3 to 4 d at 28°C before assessing growth. All experiments were performed at least three times.

### Sequence alignment, phylogenetic and gene co-expression analyses

Protein sequences from yeast Nam8 and RBP45/47 family members from the indicated plant species were obtained from the Saccharomyces Genome Database (SGD) (https://www.yeastgenome.org/) and PLAZA database V4.0 (Van Bel et al., 2018). Sequences were aligned and the phylogenetic tree generated with the software MEGA-X (Kumar et al., 2018). Protein alignment used the MUSCLE program and the tree was generated with the FastTree algorithm with default settings. Prion domain (PrD)-like regions among RBP45/47 family members were predicted by the web tool Prion-like amino acid composition (PLAAC; http://plaac.wi.mit.edu) (Lancaster et al., 2014). Co-expression analysis between *RBP45d* and *PRP39a* was carried out with ATTED II ver. 11.0 (https://atted.jp/) (Obayashi et al., 2018) and the platform ath-r.c5-0 (Obayashi et al., 2018).

### Co-immunoprecipitation (co-IP)

For co-IPs, 200 mg of plant tissue from 11-d-old seedlings for Col-0 or *RBP45d-5xc-Myc* overexpression lines was homogenized in liquid nitrogen. Total proteins were extracted in IP buffer (20 mM Tris-HCl pH 7.5, 100 mM NaCl, 10% [v/v] Glycerol, 0.2% [v/v] Triton X-100, 1 mM EDTA, 1 mM PMSF and 1x cOmplete™ EDTA-free Protease Inhibitor Cocktail [Roche, Switzerland]). Total protein extracts were then mixed with Pierce™ anti-c-Myc magnetic beads (Thermo Fisher Scientific, USA) and incubated at 4°C with gentle agitation for 1 h. Beads were collected and washed four times in washing buffer (20 mM Tris-HCl pH 7.5, 150 mM NaCl and 0.05% [v/v] Tween-20). Bound proteins were eluted with sample buffer at 95°C for 5 min. Eluted samples and input (20 µg total protein extracts) were analyzed by immunoblotting using anti-PRP39a (Agrisera, Sweden) and anti-cMyc (Santa Cruz, USA) primary antibodies.

### RNA immunoprecipitation (RIP) and RIP-seq

RIP was performed as previously described (Terzi and Simpson, 2009) with minor modifications. In brief, 11-d-old seedlings of Col-0 or *RBP45d-5xc-Myc* overexpression lines were collected. RNA and proteins were crosslinked by vacuum infiltration in 1% formaldehyde for 20 min. The crosslinking reaction was then stopped by adding glycine to a final concentration of 125 mM. After vacuum infiltration for another 10 min, the tissues were then washed four times with ddH_2_O before being homogenized in liquid nitrogen. 2.5 g of powder was resuspended in solution A (10 mM Tris-HCl pH 8.0, 10 mM MgCl_2_, 400 mM sucrose, 5 mM β-mercaptoethanol, 0.1 mM PMSF, 1x cOmplete™ EDTA-free Protease Inhibitor Cocktail, 50 U/mL SUPERase•In™ RNase Inhibitor) and filtered over two layers of Miracloth. The flow-through was then centrifuged for 20 min at 4°C at 1900*g*. The resulting pellet was resuspended in solution B (10 mM Tris-HCl pH 8.0, 10 mM MgCl_2_, 250 mM sucrose, 5 mM β-mercaptoethanol, 0.1 mM PMSF, 1.5% Triton X-100, 5x cOmplete™ EDTA-free Protease Inhibitor Cocktail, 50 U/mL SUPERase•In™ RNase Inhibitor) and centrifuged three more times for 10 min at 4°C at 12000*g*. The pellet was resuspended in solution C (10 mM Tris-HCl pH 8.0, 2 mM MgCl_2_, 1.7 M sucrose, 5 mM β-mercaptoethanol, 0.1 mM PMSF, 0.15% Triton X-100, 5x cOmplete™ EDTA-free Protease Inhibitor Cocktail, 50 U/mL SUPERase•In™ RNase Inhibitor) and centrifuged for 40 min at 4°C at 16,000*g*. The nuclei in the pellet were resuspended in solution D (50 mM Tris-HCl pH 8.0, 10 mM EDTA, 1% SDS, 0.2 mM PMSF, 10x cOmplete™ EDTA-free Protease Inhibitor Cocktail, 160 U/mL SUPERase•In™ RNase Inhibitor). Nucleic acids were fragmented with a Diagenode Bioruptor^®^ Sonicator (Diagenode, Belgium). For immunoprecipitation, anti-c-Myc magnetic beads (Thermo Fisher Scientific, USA) were first washed in RIP dilution buffer (16.7 mM Tris-HCl pH 8.0, 1.2 mM EDTA, 1.1% Triton X-100, 167 mM NaCl, 0.2 mM PMSF, 1x cOmplete™ EDTA-free Protease Inhibitor Cocktail, 160 U/mL SUPERase•In™ RNase Inhibitor), blocked in blocking buffer (1 mg/mL BSA, 0.5 mg/mL salmon sperm DNA in RIP dilution buffer), before incubation with nuclei extracts diluted ten-fold at 4°C overnight with gentle rotation. The beads were then washed three times and incubated at 70°C for 1 h to reverse crosslinking. Proteins were digested with Proteinase K and RNA was extracted with the TRIzol™ reagent (Invitrogen, USA). Contaminating genomic DNA was removed by digestion with 40 U/mL of TURBO™ DNase (Invitrogen, USA) for 30 min at room temperature. The remaining RNA was recovered with RNA Clean & Concentrator-5 (Zymo Research, USA). RNA fragments ranging from 200–1000 nucleotides in size were validated on an Agilent 2100 Bioanalyzer.

Sequencing libraries were prepared with the Illumina TruSeq stranded mRNA sample preparation kit according to the manufacturer’s protocol (Illumina, USA). In brief, purified RNA was fragmented and first-strand cDNAs were synthesized with SuperScript III reverse transcriptase (Invitrogen, USA) using dNTPs and random hexamers. Second-strand cDNAs were generated using a dUTP mix. cDNAs were then modified and ligated with Accel-NGS 2S Indexed Adapters (Swift BioScience, USA) and Accel-NGS 2S Plus DNA Library Kits (Swift BioScience, USA) according to the manufacturer’s instructions. Molecular identifiers (MIDs) of 9 nucleotides were incorporated on the i5 adapter during ligation, allowing identification of PCR duplicates. The final products were purified and enriched with 10 PCR cycles using the KAPA HiFi HotStart ReadyMix (Roche, Switzerland) and primers mix from the Swift library kit. Library concentration was determined on a BioRad QX200 Droplet Digital PCR EvaGreen supermix system (BioRad, USA) and library quality was assessed with the High Sensitivity DNA Analysis Kit (Agilent, USA). The prepared libraries were pooled for paired-end sequencing using Illumina NovaSeq at Omics Drive Co. (International plaza, Singapore) as 150-bp paired-end reads.

### Data analysis for RIP-seq

RIP-Seq reads were mapped to the Arabidopsis genome (ARAPORT11 annotation) with HISAT2 (version 2.2.2.1) with default pair-end parameters (Kim et al., 2015; Kim et al., 2019). Differential abundance of sensitive peaks was detected with ASPeak (version 2.0.1), which includes the RNA-seq data as background and the mock data as control (Kucukural et al., 2013). Default parameters were used for peak detection.

### RT-PCR and quantitative RT-PCR (RT-qPCR)

Total RNA was used for cDNA synthesis with SuperScript^TM^ III reverse transcriptase following the manufacturer’s instructions (Thermo Fisher Scientific, USA). Most primers used in RT-qPCR experiments were designed with the web-based primer design tool QuantPrime (Arvidsson et al., 2008). *PROTEIN PHOSPHATASE 2A subunit A3* (*PP2AA3*) was used as internal control for most experiments. RT-qPCR experiments were performed on a QuantStudio 12K Flex Real-Time PCR System with *Power* SYBR Green Master Mix (Thermo Fisher Scientific, USA). RT-PCR was performed with the GoTaq^®^ Green Master Mix as per manufacturer’s instructions (Promega, USA). For RIP analysis, first-strand cDNAs were synthesized with SuperScript^TM^ III reverse transcriptase (Thermo Fisher Scientific, USA) using random hexamers. RT-qPCR was done with gene-specific primers using *U3 snRNA* as internal control (Supplemental Data Set S4).

### Transcriptome deep sequencing (RNA-seq) and data analysis

Total RNA was extracted from 12-d-old seedlings grown at 22°C in LD conditions at ZT13 with the RNeasy^®^ Plant Mini Kit (QIAGEN, Germany) following the manufacturer’s instructions. Contaminating genomic DNA was removed by digestion with RNase-Free DNase Set (QIAGEN, Germany). Library preparation and data analysis were performed as described previously (Shih et al., 2019). Integrative Genomics Viewer (IGV) ver. 2.8.0 (Thorvaldsdóttir et al., 2013) was used for RNA-seq data visualization. UniProt-keyword and Gene Ontology enrichment analysis was carried out by using The Database for Annotation, Visualization and Integrated Discovery (DAVID) (Dennis et al., 2003) and the result was visualized by GO plot using R packages (Dennis et al., 2003; Walter et al., 2015).

### Accession Numbers

RNA-seq data from this publication have been submitted to the National Center for Biotechnology Information Sequence Read Archive (http://www.ncbi.nlm.nih.gov/sra) with BioProject accession numbers PRJNA735326 for RNAseq and PRJNA735425 for RIPseq. The sequence of the genes described in this study can be found at the Arabidopsis Information Resource (TAIR) website under the following accession numbers: *RBP45d* (At5g19350); *RBP45a* (At5g54900); *RBP45b* (At1g11650); *RBP45c* (At4g27000); *RBP47a* (At1g49600); *RBP47b* (At3g19130); *RBP47c* (At1g47490); *RBP47c’* (At1g47500); *U1A* (At2g47580); *U1-70K* (At3g50670); *U1C* (At4g03120); *PRP39a* (At1g04080); *ACT2* (At3g18780); *PP2AA3* (At1g13320); *FLC* (At5g10140); *FT* (At1g65480); *FLD* (At3g10390); *FLK* (At3g04610); *FCA* (At4g16280); *FY* (At5g13480); *FPA* (At2g43410); *FVE* (At2g19520); *LD* (At4g02560); *FLM* (At1g77080).

## Author Contributions

S.-L.T. and P.C. designed the experiments; P.C. performed the experiments; S.-L.T., H.-Y.H., and P.C. conducted RNA-seq data analysis; P.C. and S.-L.T. wrote the manuscript; S.-L.T. coordinated the project.

## Acknowledgments

We thank Tomonao Matsushita and Marjori Matzke for valuable discussion and Shu-Jen Chou and Ai-Ping Chen at the Genomic Technology Core Laboratory, Wen-Dar Lin at the Bioinformatics Core Laboratory of the Institute of Plant and Microbial Biology, Academia Sinica and Choun-Sea Lin, Chen-Tran Hsu and Fu-Hui Wu at the Plant Technology Core Laboratory of the Agricultural Biotechnology Research Center, Academia Sinica for technical assistance. We also thank Marjori Matzke for providing seeds of *prp39a-3*, *prp39a-4, rbp45d-1, rbp45d-2* and ST lines and Ho-Ming Chen for sharing *upf1-1*, *upf1-5* and *upf3-1* seeds. This work was supported by a grant to S.-L.T. from Academia Sinica (Grant No. AS-108-TP-L01) and Ministry of Science and Technology (Grant No. MOST 109-2311-B-001 -029 -MY3).

## Supplemental Data

**Supplemental Figure S1.** Comparison of protein sequence between Nam8 and members of the RBP45/47 family.

**Supplemental Figure S2.** RBP45d interacts with U1 snRNP components through its C terminus.

**Supplemental Figure S3.** Partial alignment of protein sequences for RBP45/45 family members from 19 species.

**Supplemental Figure S4.** Phylogenetic analysis of RBP45/47 family members.

**Supplemental Figure S5.** Generation of the *rbp45d-3* mutant by CRISPR/Cas9 genome editing.

**Supplemental Figure S6.** Quantitative analysis of flowering time for the *rbp45d-1* and *rbp45d-2* mutants in the ST background.

**Supplemental Figure S7.** Relative transcript levels of *FLC* and *COOLAIR* in *rbp45d-3* and *prp39a-1* mutants.

**Supplemental Figure S8.** Gene co-expression analysis between *RBP45d* and *PRP39a*.

**Supplemental Figure S9.** Feature analysis of DAS genes in *rbp45d-3* and *prp39a-1*.

**Supplemental Figure S10.** RBP45d regulates *PRP39a* pre-mRNA splicing.

**Supplemental Figure S11.** Overlap between RBP45d targets and differentially expressed genes (DEGs) in the *rbp45d-3* mutant relative to the ST wild-type background.

**Supplemental Figure S12.** Transcript levels of key flowering genes from the RNA-seq data.

**Supplemental Figure S13.** Temperature-responsive AS of *FLM* in *rbp45d-3* and *prp39a-1* (16°C to 28°C).

**Supplemental Figure S14.** PLAAC analysis of RBP45/47 family proteins and Nam8.

**Supplemental Table S1.** Sequence analysis for the N- and C-terminal regions of Nam8 and RBP45/47 family proteins.

**Supplemental Datasets S1.** Identifiers of the proteins used in phylogenetic analysis.

**Supplemental Datasets S2.** UniProt Keyword (KW) enrichment of DAS genes in the *rbp45d-3* and *prp39a-1* mutants.

**Supplemental Datasets S3.** List of RBP45d targets enriched by RIP-seq assay.

**Supplemental Datasets S4.** Primers used in this study.

