## Supplemental figures and table for "The U1 snRNP component RBP45d regulates temperature-responsive flowering in *Arabidopsis thaliana*"

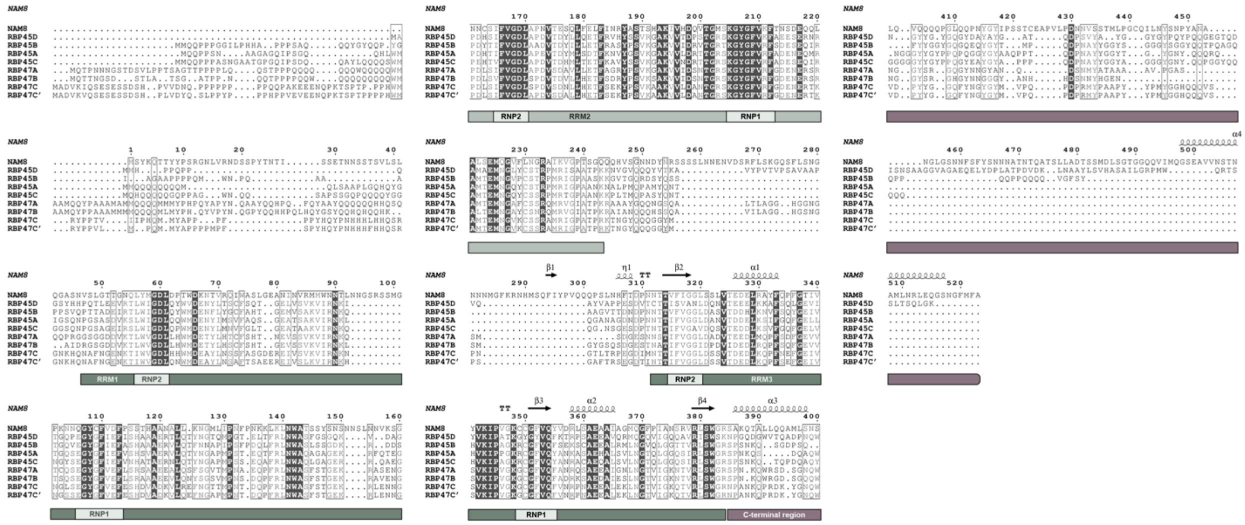

**Supplemental Figure S1.** Comparison of protein sequences between Nam8 and RBP45/47 family members. Protein sequence alignment of Arabidopsis RBP45/47 family members and yeast Nam8. The RNA recognition motifs (RRMs, sage green) and the C-terminal region (C-term, mauve) are indicated below the alignment.

(Supports Figure 1)

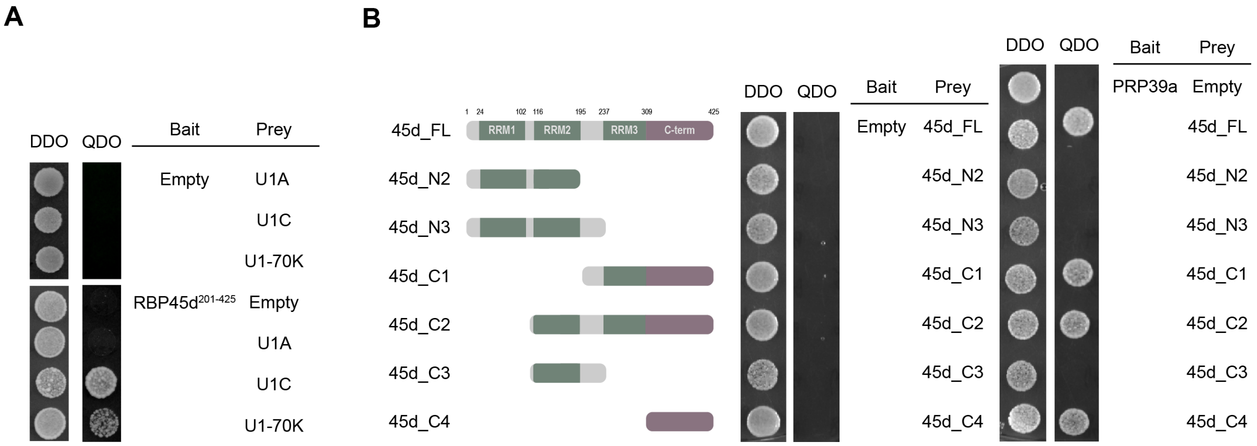


**Supplemental Figure S2.** RBP45d interacts with U1 snRNP components through its C terminus. **(A)** RBP45d interact with the two other U1 snRNP core components U1C and U1-70K in a Y2H assay. Yeast cells co-transformed with constructs expressing *RBP45d* (encoding amino acids 201–425) and *U1A*, *U1C* and *U1-70K* were diluted to an $OD_{600}$ of 0.1 and grown on synthetic defined medium lacking Leu and Trp (DDO, double dropout) for selection and SD medium lacking Leu, Trp, His and Ade (QDO, quadruple dropout) for interaction tests. Empty bait vector (Empty) was used as control. **(B)** RBP45d interacts with PRP39a through its C terminus. Yeast cells co-transformed with fragments of *RBP45d* and *PRP39a* were diluted to an $OD_{600}$ of 0.1 and grown on DDO and QDO plates. Empty bait vector (Empty) was used as control. 45d-FL, full-length RBP45d. 45d-N2, RBP45d amino acids 1–195, which include RRM1 and RRM2. 45d-N3, RBP45d amino acids 1–237. 45d-C1, RBP45d amino acids 195–425, which include RRM3 and the C terminus. 45d-C2, RBP45d amino acids 116–425. 45d-C3, RBP45d amino acids 116–237. 45d-C3, RBP45d amino acids 309–425, which include the C terminus.

(Supports Figure 1)

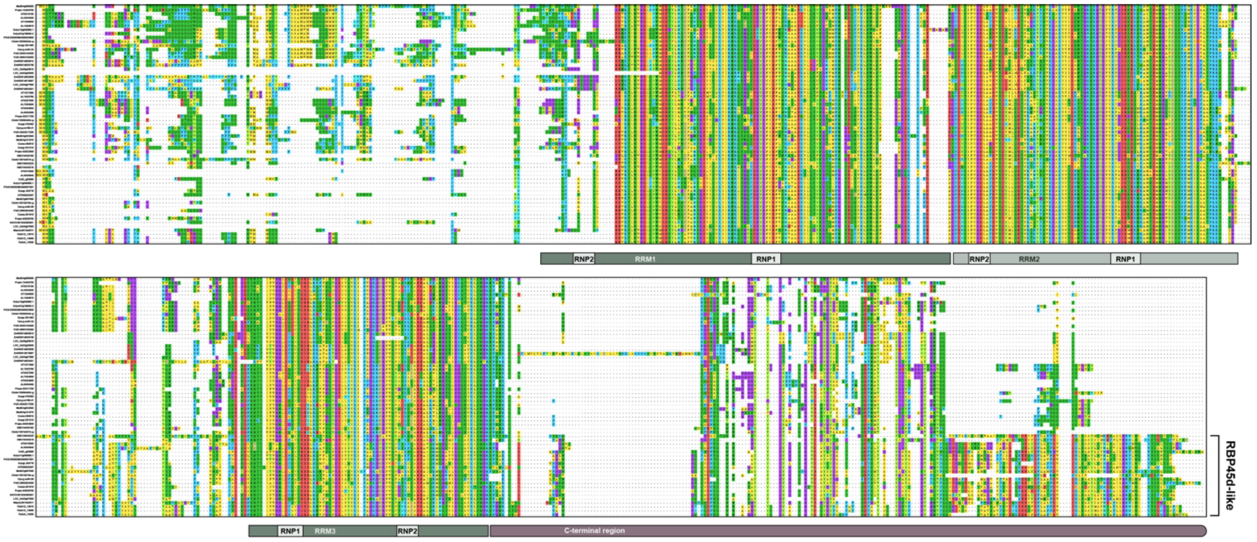


**Supplemental Figure S3.** Partial protein sequence alignment of RBP45/45 family members from 19 species. RRMs (sage green) and the long C terminus (mauve) are indicated below the alignment. All RBP45-like proteins (those with the long C terminus) were rearranged to the bottom of the alignment manually.

(Supports Figure 1)

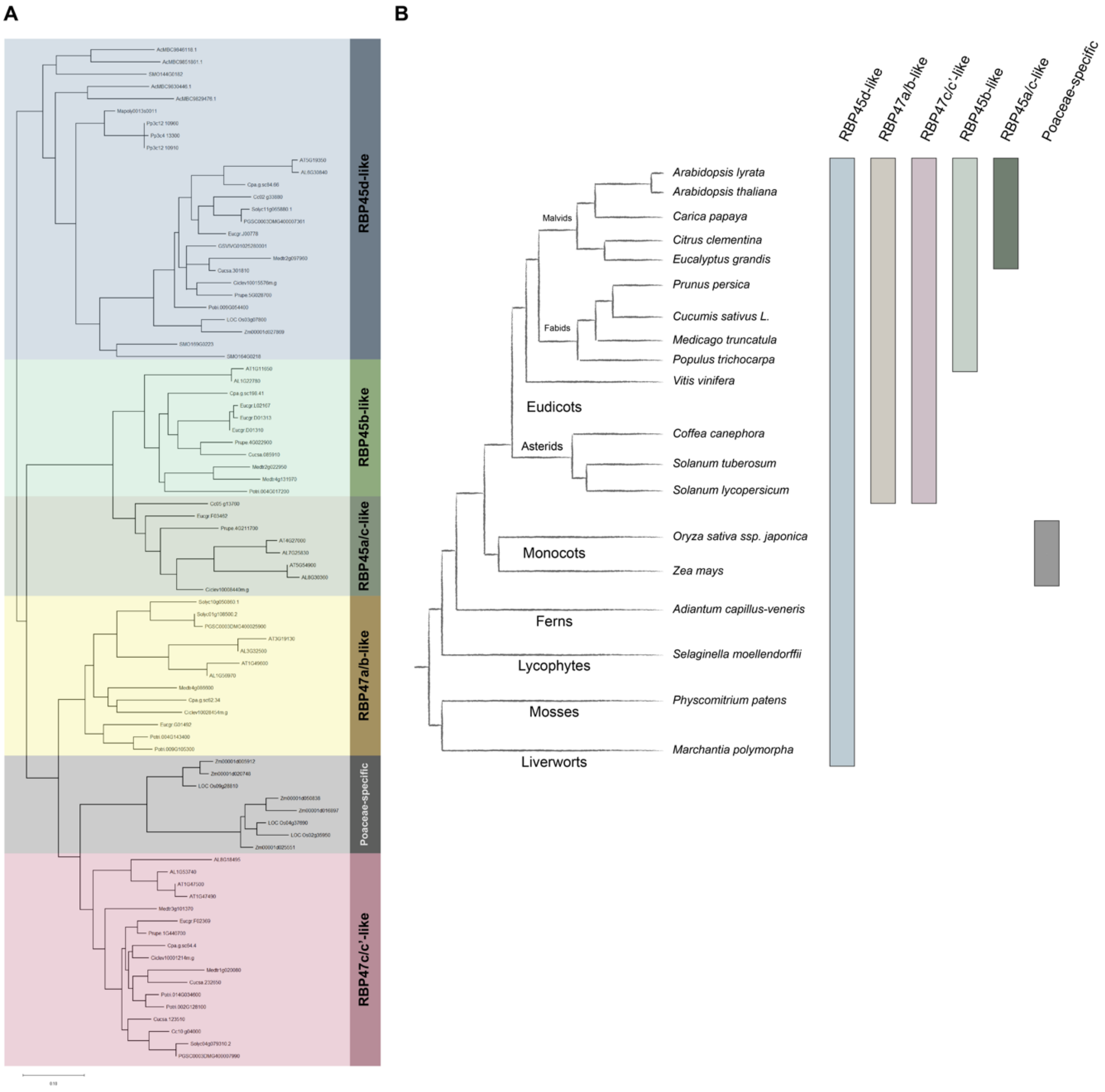


**Supplemental Figure S4.** Phylogenetic analysis of the RBP45/47 family. **(A)** Phylogenetic analysis of 84 RBP45/47 family members from 19 representative species. The tree was generated with the FastTree algorithm on the PLAZA website. The naming of groups is based on the name of their closest Arabidopsis counterpart in each clade. **(B)** Left panel, phylogenetic tree showing the evolutionary relationships between plant lineages. Right panel, the rectangle below each RBP45/47 type indicates the existence of such type in the corresponding plant lineage.

(Supports Figure 1)

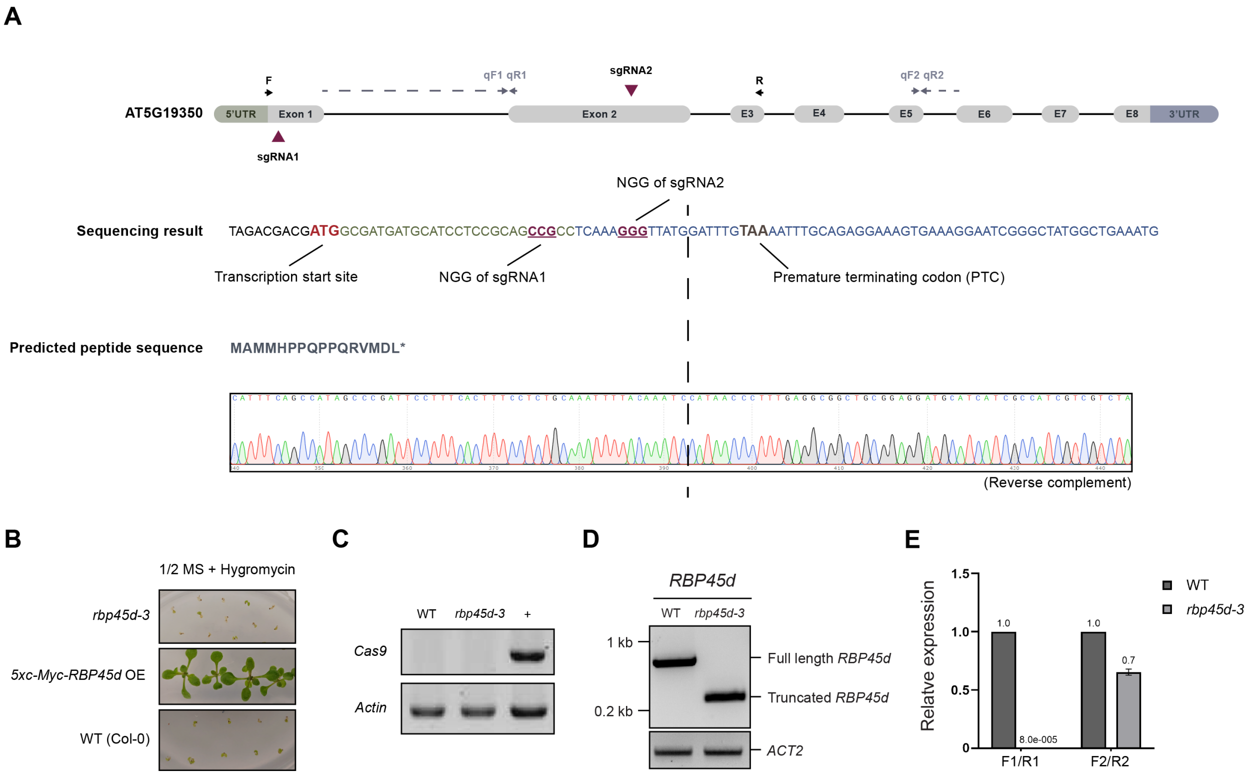


**Supplemental Figure S5.** Generation of the *rbp45d-3* mutant by CRISPR/Cas9. **(A)** Schematic representation of the *RBP45d* gene. Sequencing of *RBP45d* in the *rbp45d-3* background identified a deletion between the sites targeted by the two simple guide RNAs (sgRNA1 and 2). The predicted protein sequence translated from the edited *RBP45d* gene is also shown. The sgRNA target sites are indicated as red arrowheads. F and R, primers used for genotyping RT-PCR. qF1/qR1 and qF2/qR2, primer sets used in RT-qPCR. The removal of the Cas9+hyg cassette in *rbp45d-3* was confirmed by a survival test of seedlings grown on half-strength MS medium containing 20 µg/mL of hygromycin B Gold **(B)** and by genotyping using Cas9-specific primers **(C)**. RT-PCR **(D)** and RT-qPCR **(E)** validation of the *RBP45d* mutation in *rbp45d-3*. *ACT2* was used as internal control. Two sets of primers were used, with one designed within the deleted region (F1/R1) and one outside the deleted region (F2/R2). *PP2AA3* was used as internal control. Error bars represent standard error of the mean.

(Supports Figure 2)

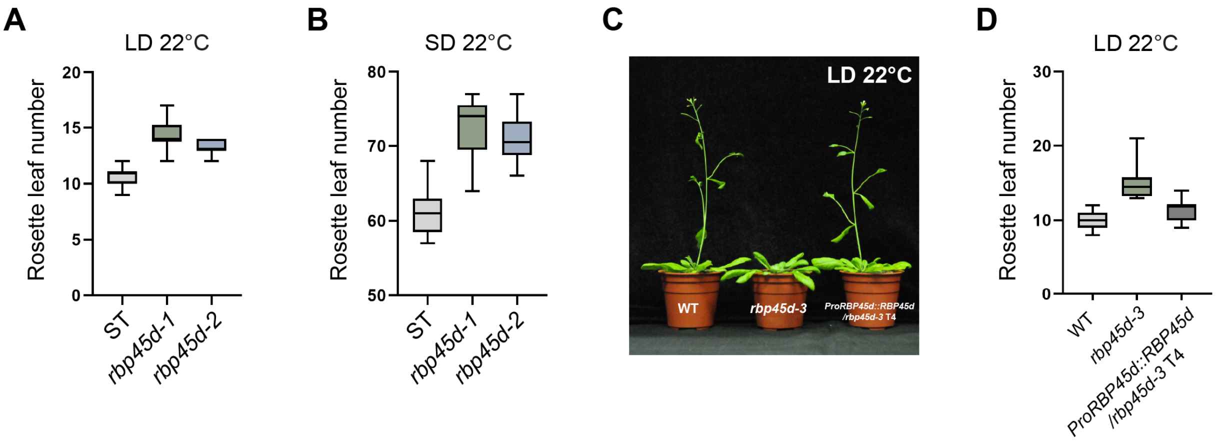


**Supplemental Figure S6.** Flowering time of other *rbp45d* alleles. Mean rosette leaf number of two other *rbp45d* alleles, *rbp45d-1* and *rbp45d-2* (Kanno et al., 2020), at bolting in LD **(A)** and SD **(B)** conditions. The *rbp45d-1* and *rbp45d-2* mutants were generated in the ST genetic background, thus the ST line was used as control. **(C)** representative photographs of WT, *rbp45d-3* and a complementation line (*rbp45d-3 RBP45dpro:RBP45d*) grown in LD conditions at 22ºC. **(D)** Mean rosette leaf number at bolting of WT (Col-0), *rbp45d-3* and the complementation line grown in LD conditions at 22ºC. At least 12 plants were scored and the experiments were performed at least three times.

(Supports Figure 2)

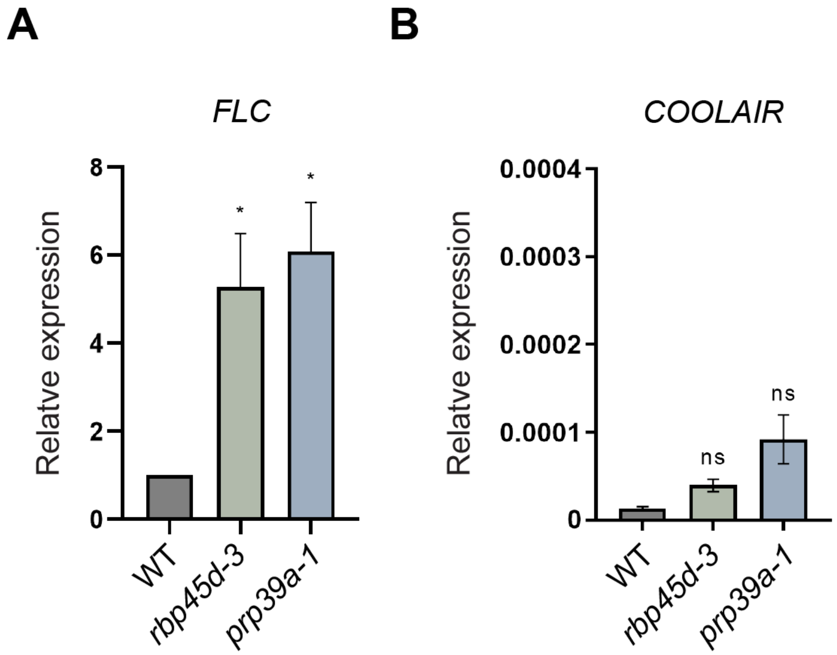


**Supplemental Figure S7.** Transcript levels of *FLC* and *COOLAIR* in the *rbp45d-3* and *prp39a-1* mutants. Error bars represent standard error of the mean. * $P < 0.05$; ns, not significant; as determined by Student's *t*-test.

(Supports Figure 2)

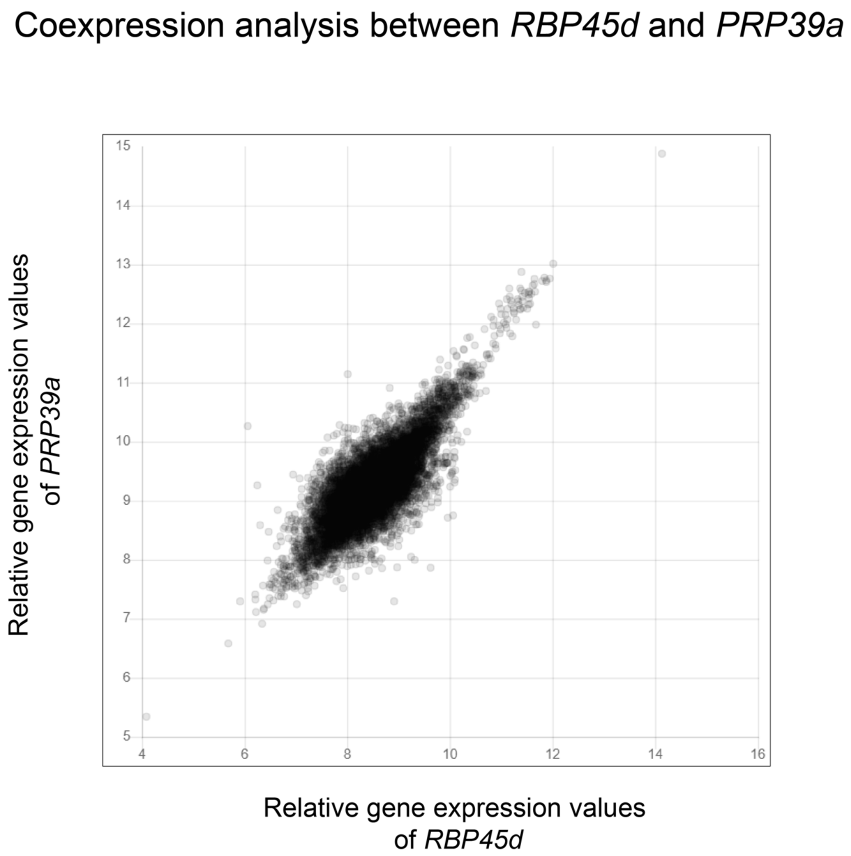


**Supplemental Figure S8.** Gene co-expression analysis between *RBP45d* and *PRP39a*. The analysis was conducted with CoexViewer on the ATTED-II website (Obayashi et al., 2018). Both axes indicate relative gene expression values ($log_2$-transformed).

(Supports Figure 3)

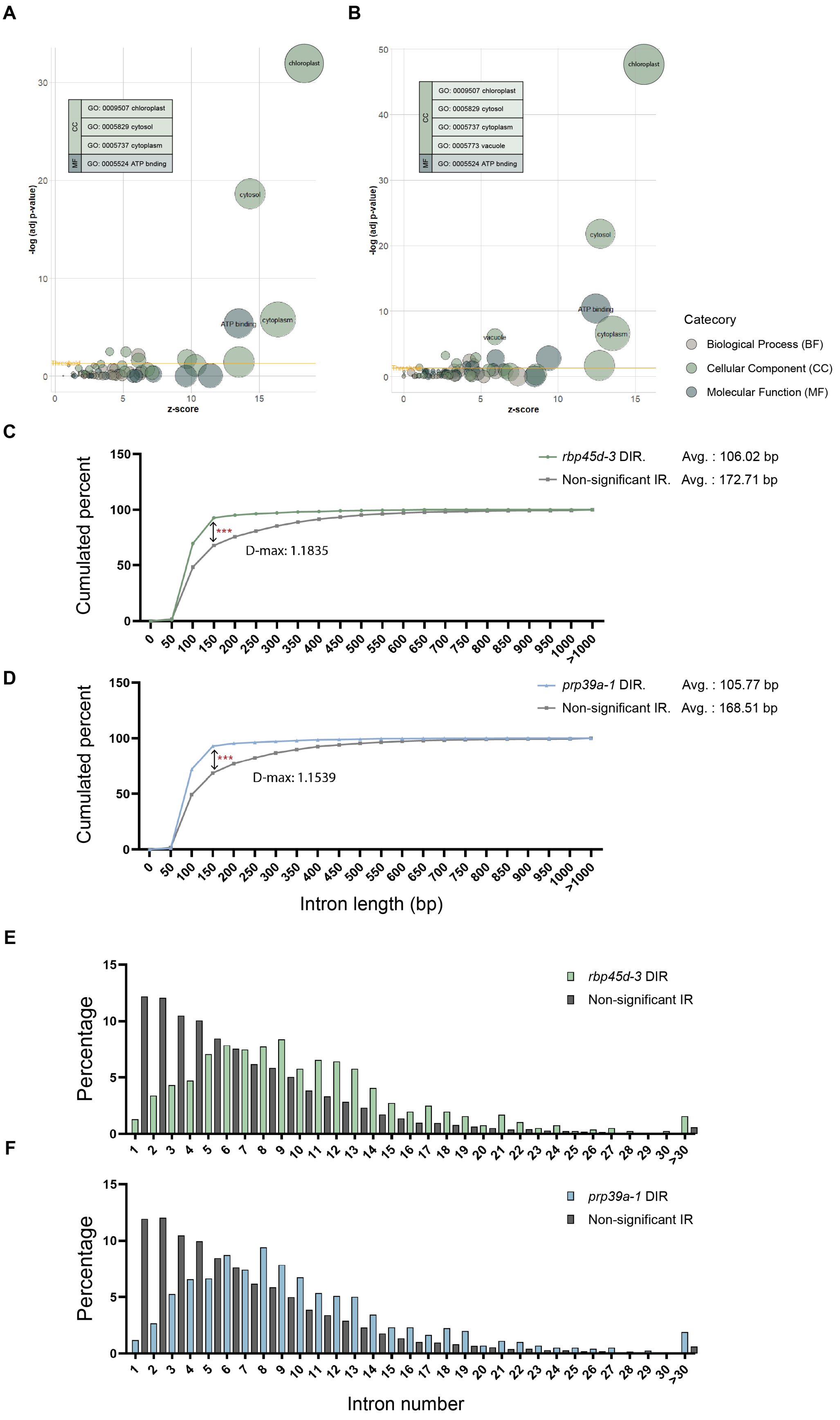
A
B
GO: 0009507 chloroplast
GO: 0005829 cytosol
GO: 0005737 cytoplasm
GO: 0005524 ATP bnding
GO: 0005773 vacuole
CC
MF
chloroplast
cytosol
cytoplasm
ATP binding
vacuole
Threshold
-log (adj p-value)
z-score
Catecory
Biological Process (BF)
Cellular Component (CC)
Molecular Function (MF)
C
rbp45d-3 DIR. Avg. : 106.02 bp
Non-significant IR. Avg. : 172.71 bp
D-max: 1.1835
***
Cumulated percent
D
prp39a-1 DIR. Avg. : 105.77 bp
Non-significant IR. Avg. : 168.51 bp
D-max: 1.1539
***
Cumulated percent
Intron length (bp)
E
rbp45d-3 DIR
Non-significant IR
Percentage
F
prp39a-1 DIR
Non-significant IR
Percentage
Intron number

**Supplemental Figure S9.** Feature analysis of DAS genes in *rbp45d-3* and *prp39a-1*. **(A** and **B)** GO analysis for DAS genes in *rbp45d-3* **(A)** and *prp39a-1* **(B)**. Each circle represents a GO term enriched from the gene sets. The names of the GO terms are shown only when the term passed the criteria (–log adjusted *p*-value) > 4; less than 75% redundancy). GO terms of different categories are indicated by different colors. **(C** and **D)** Length distribution of differentially retained introns in *rbp45d-3* **(C)** and *prp39a-1* **(D)**. Lengths of retained introns that showed no significant difference (compared to WT) in *rbp45d-3* and *prp39a-1* (non-significant IR) were used as control. A Kolmogorov-Smirnov (KS) test was used to calculate the significance of difference between the distributions. *** *P* value from KS test table < 0.001. **(E and F)** Intron number distribution of DIR genes in *rbp45d-3* **(E)** and *prp39a-1* **(F)**. Intron numbers greater than 30 were grouped into the same bin. DIR genes showing no significant difference (compared to WT) in *rbp45d-3* and *prp39a-1* (non-significant IR) were used as control.
(Supports Figure 3)

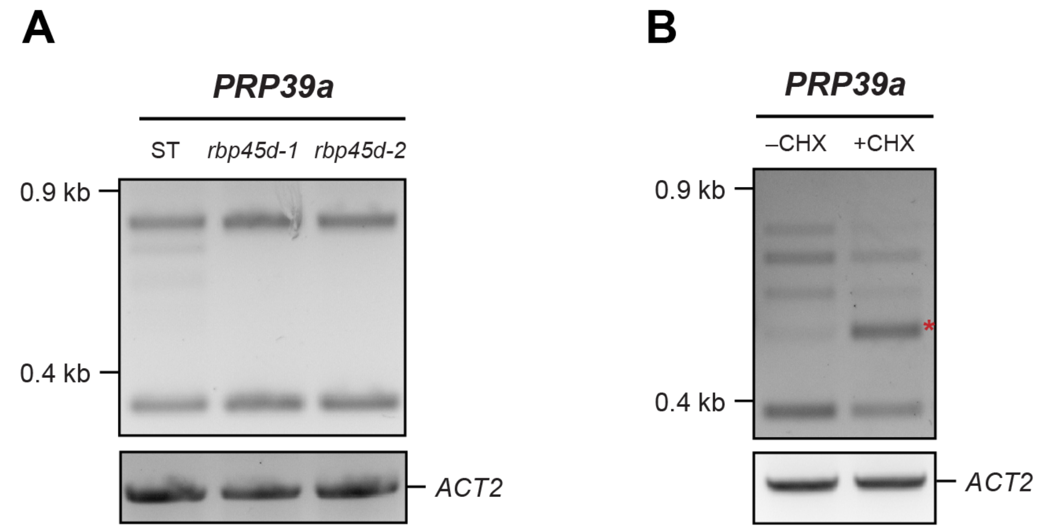


**Supplemental Figure S10.** RBP45dregulates*PRP39a* pre-mRNA splicing. **(A)** End-point RT-PCR analysis of *PRP39a* AS in ST, *rbp45d-1* and *rbp45d-2*. *ACT2* was used as internal control. **(B)** Some *PRP39a* AS isoforms are nonsense-mediated decay (NMD) targets. End-point RT-PCR analysis of *PRP39a* AS in plants treated with cycloheximide (+CHX) or with DMSO (–CHX) for 4 h. *ACT2* was used as internal control.

(Supports Figure 4)

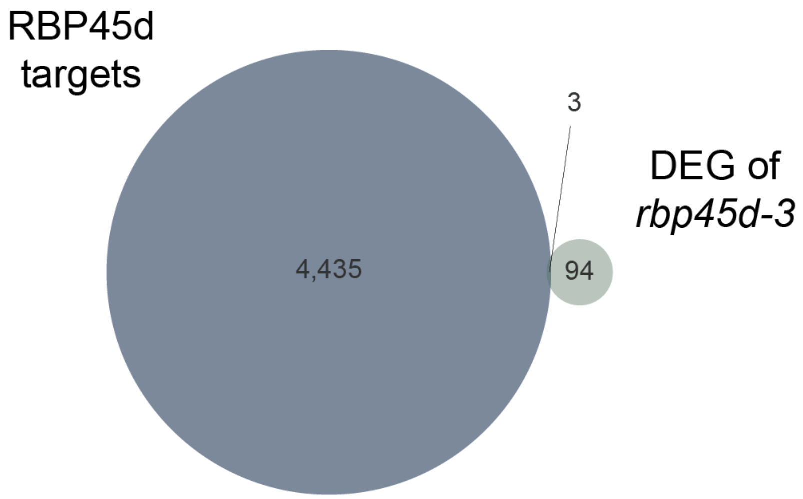


**Supplemental Figure S11.** Venn diagram of the overlap between RBP45d targets enriched determined by ASPeak and DEGs in *rbp45d-3*.
(Supports Figure 5)

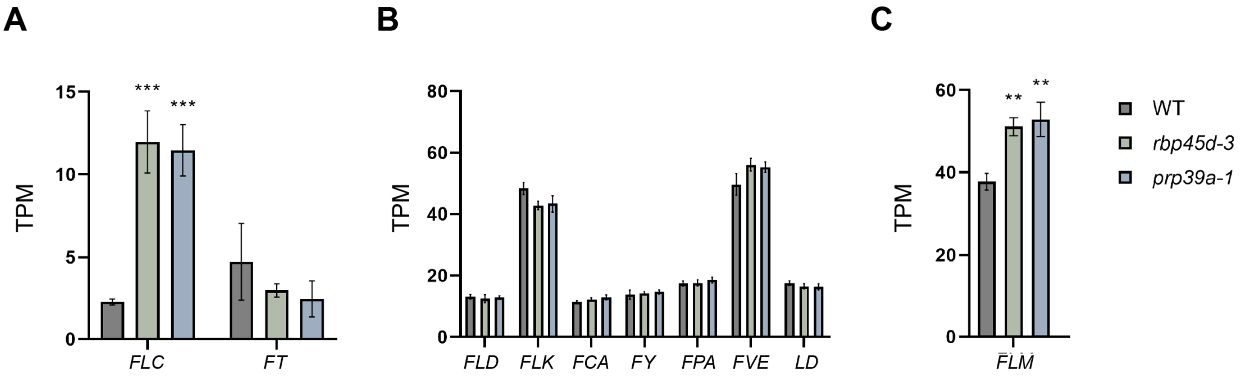


**Supplemental Figure S12.** Transcript levels of key flowering genes from RNA-seq. **(A)** Transcript levels of *FLC* and *FT* in WT, *rbp45d-3* and *prp39a-1*. **(B)** Transcript levels of key genes in autonomous pathway including *FLD*, *FLK*, *FCA*, *FY*, *FPA*, *FVE* and *LD* in WT*, rbp45d-3* and *prp39a-1*. **(C)** Transcript levels of *FLM* in WT, *rbp45d-3* and *prp39a-1*. All transcript levels are shown as Transcripts Per Kilobase Million (TPM) and the error bars represent standard deviation ($n = 3$). ** $P < 0.01$; *** $P < 0.001$ as determined by Student's *t*-test.

(Supports Figure 6)

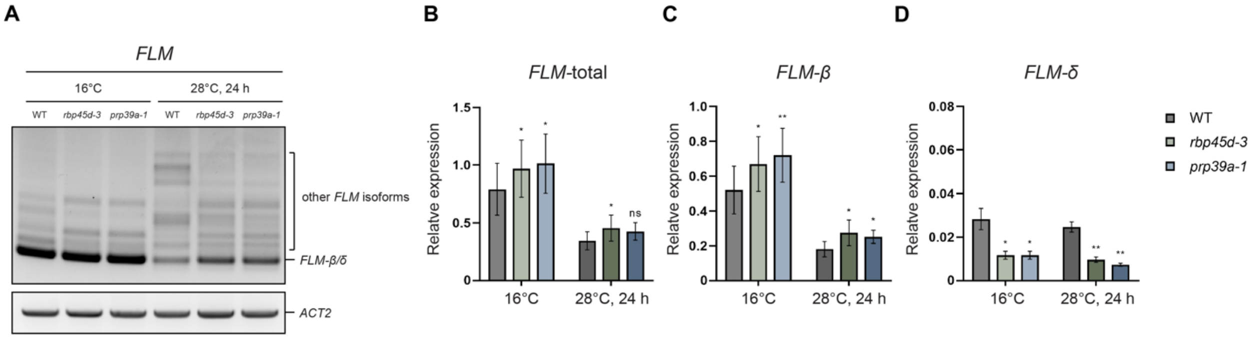


**Supplemental Figure S13.** Expression level of *FLM* isoforms under raising temperature. **(A)** End-point RT-PCR analysis of *FLM* AS isoforms in WT, *rbp45d-3* and *prp39a-1* grown at 16°C or shifted from 16°C to 28°C for 24 h. *ACT2* was used as internal control. **(B-D)** RT-qPCR of bulk *FLM* transcript levels and of its two major isoforms *FLM-β* and *FLM-δ* in WT, *rbp45d-3* and *prp39a-1 1* grown at 16°C or shifted from 16°C to 28°C for 24 h. Error bars represent standard error of the mean. * $P < 0.05$; ** $P < 0.01$ as determined by Student's *t*-test. *PP2AA3* was used as internal control.

(Supports Figure 7)

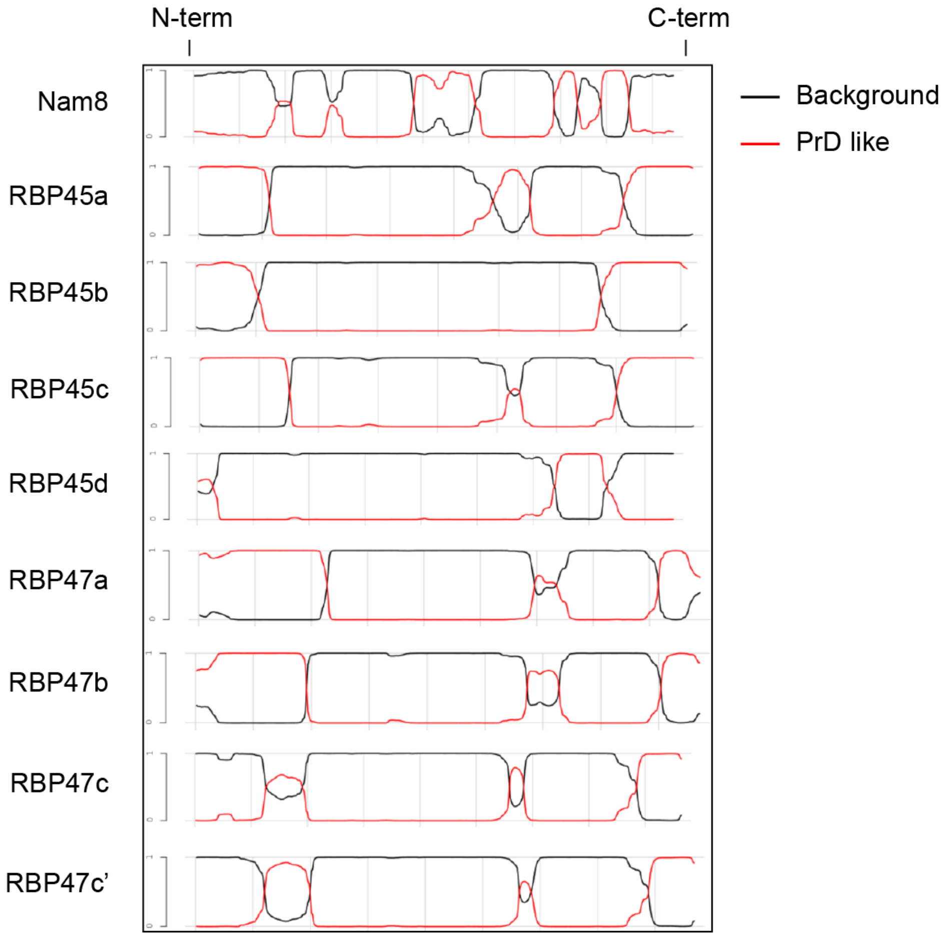


**Supplemental Figure S14.** PLAAC analysis of RBP45/47 family proteins and Nam8. (Supports the Discussion)

**Supplemental Table 1.** Amino acid sequence analysis for the N-terminal **(A)** and C-terminal **(B)** regions of Nam8 and RBP45/47 family proteins.

**(A)** N terminus analysis of Nam8 and RBP45/47 family proteins. Gln(Q); Asn (N); Tyr (Y); Gly (G).

| | N-term length | Q | N | Y | G | QNYG% |
|---|---|---|---|---|---|---|
| Nam8 | 47 | 2 | 6 | 4 | 2 | 30% |
| RBP45a | 53 | 17 | 2 | 1 | 6 | 49% |
| RBP45b | 55 | 10 | 1 | 3 | 5 | 35% |
| RBP45c | 73 | 24 | 3 | 2 | 10 | 53% |
| RBP45d | 17 | 2 | 0 | 1 | 1 | 24% |
| RBP47a | 112 | 37 | 4 | 7 | 4 | 46% |
| RBP47b | 101 | 30 | 3 | 6 | 3 | 42% |
| RBP47c | 94 | 9 | 3 | 4 | 1 | 18% |
| RBP47c’ | 96 | 7 | 2 | 6 | 1 | 17% |

**(B)** C terminus analysis of Nam8 and RBP45/47 family proteins.

| | C-term length | Q | N | Y | G | QNYG% |
|---|---|---|---|---|---|---|
| Nam8 | 139 | 16 | 18 | 5 | 11 | 36% |
| RBP45a | 56 | 11 | 4 | 10 | 11 | 64% |
| RBP45b | 73 | 17 | 2 | 11 | 15 | 61% |
| RBP45c | 66 | 13 | 3 | 13 | 16 | 68% |
| RBP45d | 116 | 12 | 4 | 13 | 17 | 39% |
| RBP47a | 47 | 4 | 7 | 4 | 7 | 47% |
| RBP47b | 43 | 3 | 8 | 4 | 10 | 58% |
| RBP47c | 57 | 7 | 4 | 10 | 7 | 49% |
| RBP47c’ | 57 | 7 | 4 | 10 | 7 | 49% |
